# Yeast *de novo* proteins integrate into cellular systems using ancient protein targeting and degradation pathways

**DOI:** 10.1101/2024.08.28.610198

**Authors:** Carly J. Houghton, Nelson Castilho Coelho, Annette Chiang, Aaron Wacholder, Stefanie Hedayati, Alison Guyer, Matayo Wankiiri-Hale, Hussain Raza, Nathan D. Lord, Saurin B. Parikh, Nejla Ozbaki-Yagan, John Iannotta, Alexis Berger, Matthew L. Wohlever, Anne-Ruxandra Carvunis, Allyson F. O’Donnell

## Abstract

Recent evidence demonstrates that eukaryotic genomes encode thousands of evolutionarily novel proteins that originate *de novo* from non-coding DNA and can contribute to species-specific adaptations. Yet, it remains unclear how these incipient proteins—whose sequences are entirely new to nature—navigate the cellular environment to bring about phenotypic change. Here, we conduct a systematic *in vivo* investigation of yeast *de novo* proteins with enhanced growth phenotypes, revealing the early stages of cellular integration. We find that these proteins are strongly enriched at the endoplasmic reticulum (ER) relative to conserved proteins, and that they integrate into cellular systems through conserved membrane targeting, trafficking, and degradation pathways. Despite having unrelated sequences, ER-localized *de novo* proteins share a common molecular signature: a C-terminal transmembrane domain that likely enables recognition by conserved post-translational ER insertion pathways. After insertion, ER-localized *de novo* proteins traffic from the ER and their homeostasis is regulated by conserved proteasomal and vacuolar degradation pathways. Our findings demonstrate that ancient targeting and degradation pathways can accommodate young *de novo* proteins sharing a convergent molecular signature. These pathways may act as selective filters, biasing which young *de novo* proteins persist.

**Significance Statement:** Deeply conserved genes shape the core structure and function of cells, but novel genes are key drivers of biodiversity. Novel genes can arise by divergence from ancient genes, or *de novo* from non-genic DNA. For *de novo* genes to develop novel functions, it is necessary for them to be processed by the cell so that their protein levels and localization are regulated. However, little is known about how novel *de novo* genes obtain the capacity to engage the cellular machinery needed for this regulation. Here, we experimentally assess how a set of endoplasmic reticulum (ER)-localized proteins encoded by recently-evolved *de novo* genes are localized and degraded in yeast cells. We discover that, despite having entirely unique amino acid sequences, these proteins share biochemical signatures allowing them to engage the same ancient cellular machinery and localize to the ER membrane. Interestingly though, this machinery is not the one that targets most ancient proteins to the ER. These results indicate that even recently emerged proteins without an extensive period of evolutionary adaptation can be recognized by specific ancient cellular pathways, facilitating their localization and homeostasis.

## Introduction

New protein-coding genes can evolve *de novo* from sequences that were previously non-genic. Once considered rare, *de novo* gene birth is now recognized as a major force of molecular innovation and adaptation across species (1–4). However, the mechanisms by which these *de novo* proteins integrate into cellular systems remain poorly understood (4, 5). Ancient proteins have been coevolving for millions of years with the systems that help them fold correctly, localize to their appropriate compartments, and regulate their homeostasis (6–9). How are *de novo* proteins (DNPs) — naïve to these systems — recognized and processed by the cell such that their expression is not only tolerated, but potentially beneficial (**Fig 1A**)?

**Figure 1.**
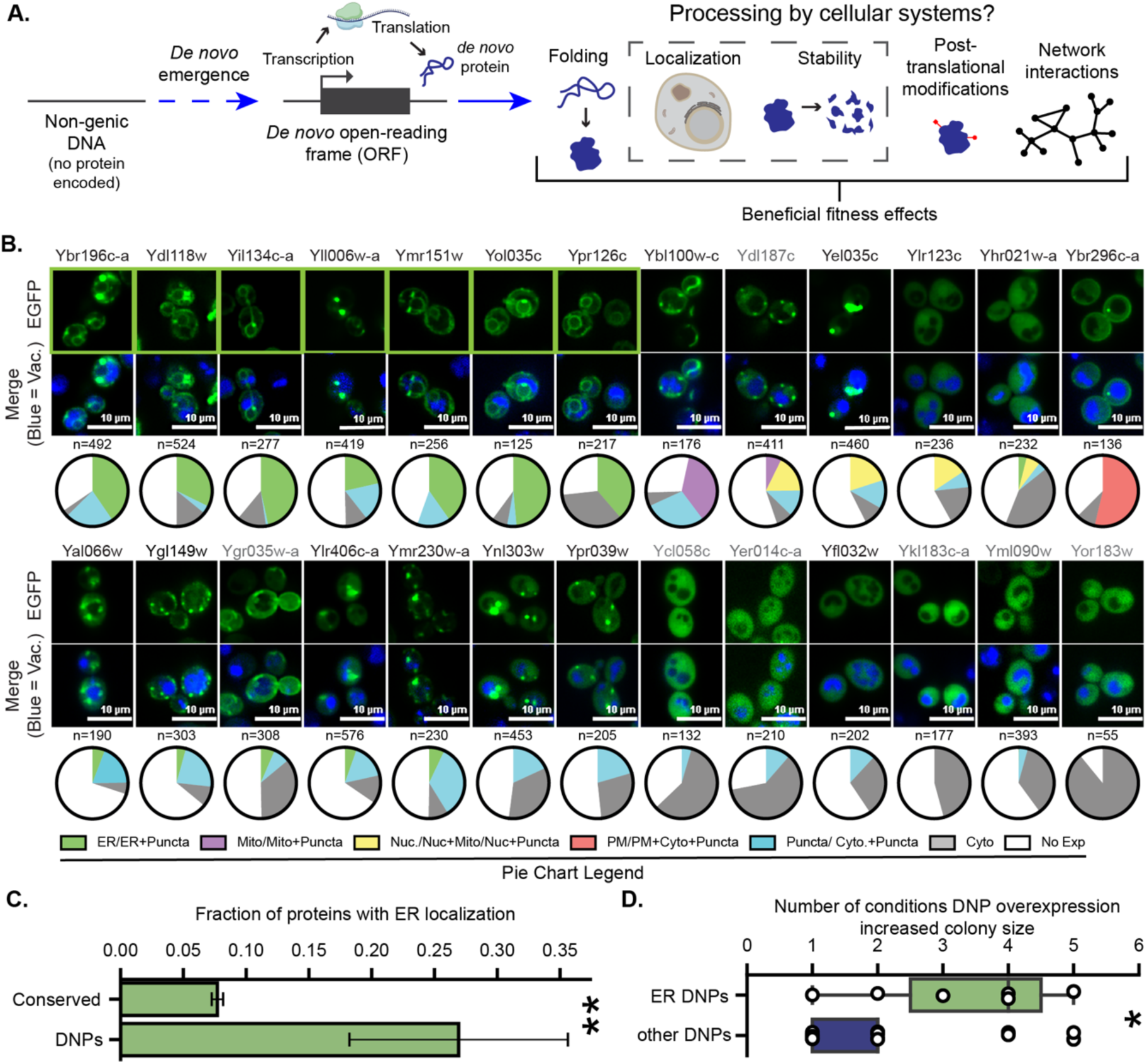
DNPs preferentially localize at the ER. **A.** Schematic representation of DNP emergence and the various processing by cellular systems that ultimately gives rise to their beneficial fitness effects. DNPs are encoded by *de novo* ORFs which are created by mutations within non-genic DNA. The specific processes and mechanisms that DNPs use to integrate into the existing cellular systems remain unknown. The processes of localization and stability (boxed by dashed outline) are the focus of the current study. **B.** Seven out of 26 DNPs localize to the ER. Fluorescence microscopy of cells induced to express DNP-EGFP fusions from a 2µ plasmid are shown. Images of ER-localized DNPs are outlined in green. Cells were stained with CMAC blue to mark the vacuoles. The distributions of localizations observed across individual cells (n) are provided as pie-chart graphs below each image. Mito = mitochondria, Nuc = nucleus, PM = plasma membrane, Cyto = cytoplasm, No Exp = no expression. The DNPs highlighted in grey correspond to those not presenting a band at the expected size on immunoblotting analysis as shown on Figure S1B. **C.** DNPs are enriched in the ER relative to conserved proteins. The fraction of all DNP-EGFP fusions in this study (n=26) that localize to the ER upon induced expression *vs* the fraction of the whole *S. cerevisiae* proteome fused C-terminally to GFP and natively expressed (n=3488, GFP collection (16)) that stably localize to the ER are compared. Error bars indicate the standard error of the proportion. **: Fisher exact test, odds ratio= 4.4, p= 3.1 x 10^-3^. **D.** ER-localized DNPs increased colony size in more conditions than other DNPs. The distribution of the number of experimental conditions in which ER-localized and other DNPs were found to be beneficial when overexpressed by Vakirlis and colleagues (12) is presented. *: Mann Whitney U test, p= 0.025, n= 26.

We can consider several hypotheses: 1) The recognition mechanisms by cellular processing systems could be sufficiently flexible that even newly-evolved proteins can make use of the full suite of pathways available to ancient proteins for folding, localization, and degradation; 2) Only a subset of standard pathways might be readily accessible to newly evolved proteins, and these specific pathways would then have a disproportionate role in facilitating the evolution of new proteins; 3) Young proteins might initially be processed by non-standard pathways before eventually evolving to use standard pathways optimally; 4) Young proteins, having not had sufficient time to evolve proper signals, could fold and localize on their own without the aid of other proteins (10, 11).

Here, we sought to better understand cellular processing of potentially adaptive DNPs through an investigation of the targeting and degradation mechanisms for a collection of *S. cerevisiae* proteins that emerged *de novo* in the *Saccharomyces* lineage. We focus specifically on a set of proteins that show a growth-enhancing phenotype upon overexpression in stress conditions, as this suggests the potential for the protein to provide an adaptive benefit to the organism. We find that growth-enhancing young DNPs preferentially localize to membrane-bound compartments, with a strong enrichment at the ER relative to the conserved proteome. We then show that this ER-targeting depends on the ancient post-translational targeting machinery in the (<u>G</u>uided <u>E</u>ntry of <u>T</u>ail-anchored proteins (GET) and <u>SRP</u>-i<u>ND</u>ependent (SND) pathways, while homeostasis of these young DNPs at the ER is regulated by the conserved <u>ER</u>-<u>a</u>ssociated <u>d</u>egradation pathway (ERAD). Engagement with the GET and SND pathways appears to be enabled by predicted C-terminal TMDs that are found in all ER-localized DNPs we identified. In contrast, young DNPs appear to lack signals enabling recognition by the co-translational SRP (signal recognition particle) pathway, the most common ER-targeting pathway among conserved ER-localized genes. These results support a model by which DNPs are recognized by standard pathways that also process conserved proteins rather than making use of DNP-specific pathways or self-targeting. However, DNPs do not appear to use the full suite of pathways available to conserved proteins; rather, some pathways are especially suited to DNPs, and this is facilitated by recognition of simple molecular signals like a C-terminal TMD. Over evolutionary timescales, these ancient pathways that are disproportionately involved in processing DNPs may play a particularly influential role in shaping their fate.

## Results

### Recently-evolved *de novo* proteins with increased growth phenotypes preferentially localize to the ER

Our aim was to understand the targeting and degradation mechanisms used by potentially adaptive *de novo* proteins (DNPs). Towards this end, we considered a set of 28 annotated yeast proteins of recent *de novo* origin that caused increased colony size relative to wildtype when overexpressed in at least one nutrient stress condition (i.e. varying nitrogen and/or carbon source composition; **Supplemental Table 1**). We previously showed that these proteins emerged in the *Saccharomyces* lineage from non-coding sequences (12). Though these proteins are poorly conserved in *Saccharomyces* and generally do not show strong loss-of-function phenotypes, the increased colony size upon overexpression of these genes suggests a potential for these proteins to be useful in future adaptation.

We first assessed where these 28 DNPs localize. We were able to successfully make C-terminal, EGFP fusions for 26 of 28 DNPs and expressed these from plasmids under the control of an inducible promoter (13–15). Microscopy revealed that most proteins tested (18/26) localized to membrane compartments (ER, mitochondria, nucleus, and plasma membrane) in at least some cells (**Fig 1B; Supplemental Table 1**). The ER was overrepresented among these compartments (**Fig 1B and Fig S1A**). Seven out of twenty-six DNPs (27%) showed enriched ER localization in contrast to only ∼ 8% of the ancient yeast proteome (Fisher exact test, odds ratio= 4.4, p= 3.1 x 10^-3^; **Fig 1C**) (16). In sum, we find that potentially adaptive DNPs localize to a wide variety of cellular compartments, yet they are enriched in the ER.

In some cells, DNPs displayed diffuse, punctate, or no measurable fluorescence (**Fig 1B**). This could be due to plasmid loss, reduced transcript stability, reduced translation efficiency, or poor protein stability. Immunoblotting revealed a broad distribution of protein abundances and breakdown products (**Fig S1B**), suggesting that DNPs are highly susceptible to degradation.

Given the unexpectedly high number of DNPs at the ER, we compared the phenotypic impacts of overexpressing ER-localized DNPs to those that do not localize to the ER. ER-localized DNPs provided growth increases across a broader array of nutrient stress conditions (12) than other DNPs (Mann Whitney U test, p= 0.025, n= 26; **Fig 1D**). These results reveal a strong association between ER localization and DNPs whose overexpression improves growth in yeast. Since these results suggest an outsized role for the ER in mediating the emergence of adaptive DNPs, we decided to focus our study on defining the degradation and targeting mechanisms used by ER-localized DNPs, assessing in particular how these proteins can be targeted to the ER despite possessing novel sequences that have had minimal opportunity to co-evolve with the cellular machinery.

### ER-localized DNPs possess C-terminal transmembrane domains

First, we searched for potential molecular determinants within the amino acid sequences of DNPs that may distinguish the ER-localized proteins from the others. All 26 DNPs are shorter than 200 amino acids; 22/26 are under 150 amino acids, and 10/26 are shorter than 100 amino acids, which is a common threshold for microprotein classification (17). Most DNPs (21/26), whether they are ER localized or not, are predicted to encode TMDs by both Phobius (18) and TMHMM (19) (**Fig S2**) (12). Structural predictions by Alpha-Fold2 (20) were consistent with these predictions revealing widespread alpha helices throughout these proteins (**Fig S3**). Thus, all the ER-localized DNPs display the biophysical potential to integrate into membranes due to the presence of predicted TMDs, but this feature alone cannot explain their specific ER localization as TMDs are also present in DNPs localizing elsewhere.

We therefore asked whether the protein sequences of ER-localized DNPs possessed unique biochemical properties distinguishing them from other DNPs. A majority of conserved proteins that localize to the ER do so co-translationally through the signal recognition particle (SRP) pathway, which recognizes a signal peptide (21, 22). However, no signal sequences were predicted in the ER-localized DNPs by TargetP 2.0 (23) or SignalP 6.0 (24). Alternatively, ER proteins can be synthesized in the cytoplasm and then post-translationally inserted in the ER using the SND and/or GET pathways (25). These pathways recognize TMDs in the middle or the C-terminus of proteins, respectively (25). In addition, membrane proteins targeted to the ER tend to have more hydrophobic TMDs and less positively charged C-tail extensions (CTEs) than proteins targeted to alternative membranes like the mitochondria (26, 27). To look for potential molecular determinants of ER localization in DNP sequences, we measured the length, hydrophobicity, and positive charge in different sequence regions from all DNPs with predicted TMDs. We considered separately their full-length sequences, regions corresponding to their N or C terminal ends, or their predicted TMDs (**Fig 2A**).

**Figure 2.**
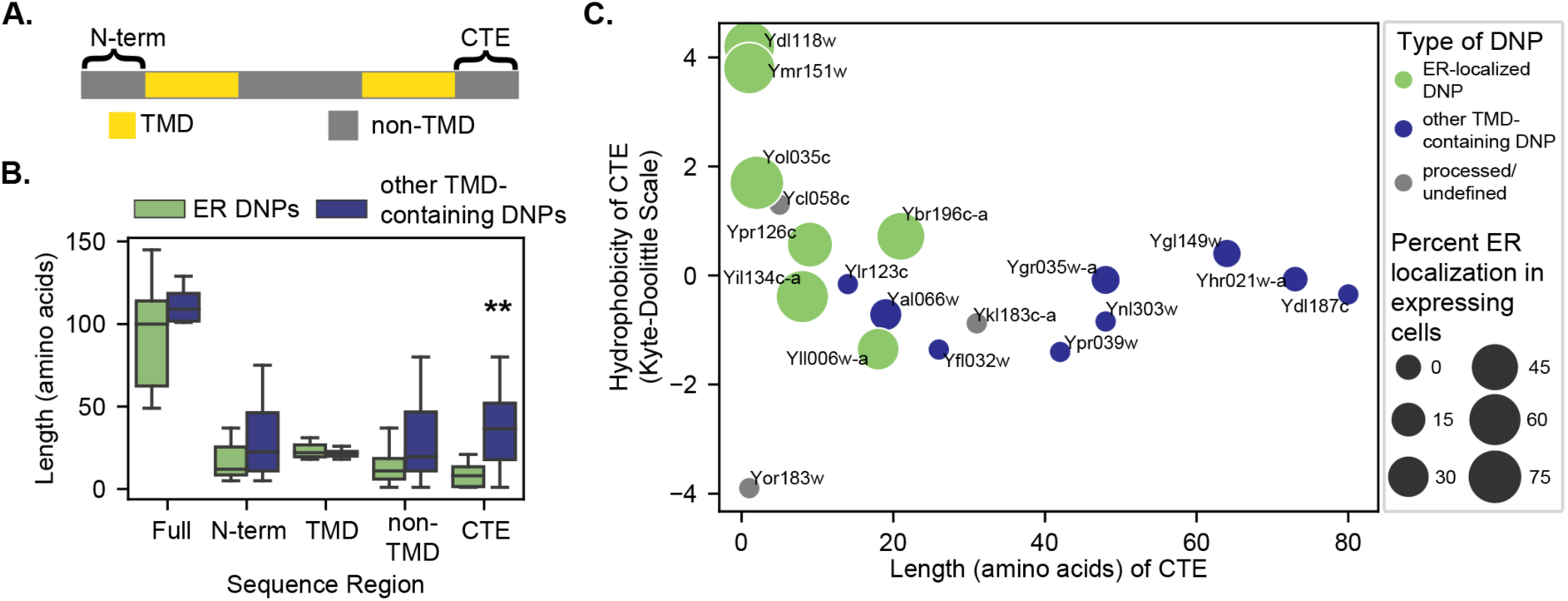
ER-localized DNPs have C-terminal TMDs. **A.** Cartoon diagram of a TMD-containing DNP sequence. Each sequence was separated into regions based on TMD predictions by either Phobius (18) or TMHMM (19) in Fig S4. These regions included 1) the N term: all residues from the start of the sequence to the start of a predicted TMD; 2) TMD: regions predicted to be TMDs; 3) non-TMD: regions including all residues not predicted to be in a TMD; and 4) the CTE: all residues following the most distal TMD in the sequence. **B.** ER-localized DNPs have significantly shorter CTEs than other TMD-containing DNPs. Length in amino acid residues for different sequence regions in ER-localized and other TMD-containing DNPs is plotted (Mann-Whitney U test. *: p<0.05; Full=Full sequence. For Full, N-term, and CTE, n=19. For TMD, n=30. For non-TMD n=49. **C.** Hydrophobicity versus length in amino acid residues for the CTEs of all TMD containing DNPs. Processed/undefined: DNPs that showed some evidence of breakdown in western blots (see Fig S1b). The size of the dots represents the percent of expressing cells that had ER localization in Figure 1B. Corresponding systematic gene names from *Saccharomyces* Genome Database shown next to each dot.

We found that all three features cleanly distinguished ER-localized DNPs from the other TMD-containing DNPs. Consistent with a post-translational ER-insertion mechanism, ER-localized DNPs were predicted to have TMDs near their C-terminus, with short CTEs less than 26 amino acids (26, 28) beyond the last predicted TMD (**Fig S4**). The predicted CTEs of ER-localized DNPs were significantly shorter than those of other TMD-containing DNPs (Mann-Whitney U test, p = 0.036, effect size r = Z/✓N = 0.48; **Fig 2B, S4A**). In contrast, the lengths of ER-localized DNPs, as well as the lengths of their N-termini or TMDs, were statistically indistinguishable from those of other TMD-containing DNPs (**Fig 2B, S4A**). In addition to being shorter, the CTEs after the final TMD of ER-localized DNPs were less positively charged and more hydrophobic than those of other TMD-containing DNPs (**Fig 2C, Fig S4B-E**). The relative paucity of positively charged residues agrees with previous work on distinguishing characteristics of post-translationally targeted ER tail-anchored (TA) proteins in yeast (27).

Upon closer inspection of the predicted structures (**Fig S3**), it becomes apparent that while several DNPs have unstructured CTEs, the CTEs of all ER-localized DNPs end with either just a few hydrophobic residues after a TMD or an alpha-helical motif. Therefore, given the uncertainty in TMD boundary predictions (29) (**Fig S2**), some of the residues in the predicted CTEs of ER-localized DNPs could actually be part of the C-terminal TMDs. If so, the true TMDs of ER-localized DNPs would be even closer to their C termini than suggested by sequence-based predictions—which are already very short and highly compatible with post-translational insertion mechanisms (28) (**Fig 2C**, **Fig S4F**).

Together our analyses show that all ER-localized DNPs are predicted to contain TMDs near their C-termini and lack predicted signal peptides. These molecular signatures are consistent with post-translational targeting of ER-localized DNPs via the GET and/or SND pathways, rather than the co-translational insertion via the SRP-dependent pathway, which recognizes signal peptides and is the most common ER insertion route among ancient proteins (21). We thus aimed to assess experimentally whether the ER localization of these DNPs is dependent on 1) their C-terminal, TMD-containing regions and 2) the GET and/or SND pathways.

### The C-terminal region of a model DNP is sufficient for ER targeting

We initially focused on the ER-localized DNP Ybr196c-a to determine if its targeting depended on its C-terminal region. This protein is a good model for ER targeting of young proteins as we previously showed that it is a bona-fide integral ER membrane protein (12). Ybr196c-a has one of the longest predicted CTEs among all ER-localized DNPs (26 residues; **Fig S2**). We suspected that this predicted CTE might mask a second TMD that failed to be predicted since the C-terminus of Ybr196c-a was predicted to be alpha-helical (**Fig S3**). Evolutionary analyses of the Ybr196c-a locus across budding yeast species are also consistent with a second TMD (12), which is predicted in the homologous region of Ybr196c-a’s orthologue in *S. paradoxus* (12).

To map the sequence determinants that dictate Ybr196c-a’s ER localization, we assessed the localization of the full-length and two discrete fragments of each orthologous Ybr196c-a sequence from *S. cerevisiae* and *S. paradoxus* spanning from: 1) the N-terminus to the end of the *S. cerevisiae*’s predicted TMD (Ybr196c-a 1.1), and 2) the end of the *S. cerevisiae*’s predicted TMD to the C-terminus of either species’ sequences (Ybr196c-a 1.2) (**Fig 3A**). We integrated the full-length and partial sequences (1.1 and 1.2) fused with mNG into the *S. cerevisiae* genome under the control of an inducible promoter (15). The full-length Ybr196c-a of both *S. cerevisiae* and *S. paradoxus* localized to the ER in 100% of cells (**Fig 3B**, left). In both species, the N-terminal sequence, Ybr196c-a 1.1, localized to the mitochondria, while the C-terminal sequence, Ybr196c-a 1.2, localized to the ER (**Fig 3B**, right). These results suggest that the CTE of Ybr196c-a in *S. cerevisiae* contains a TMD that was missed by computational prediction yet is sufficient for integration into the ER membrane (12). The fact that the C-terminal region of our model ER-localized DNP dictates its targeting to the ER is consistent with known mechanisms of post-translational insertion for conserved proteins by the GET pathway (25).

**Figure 3.**
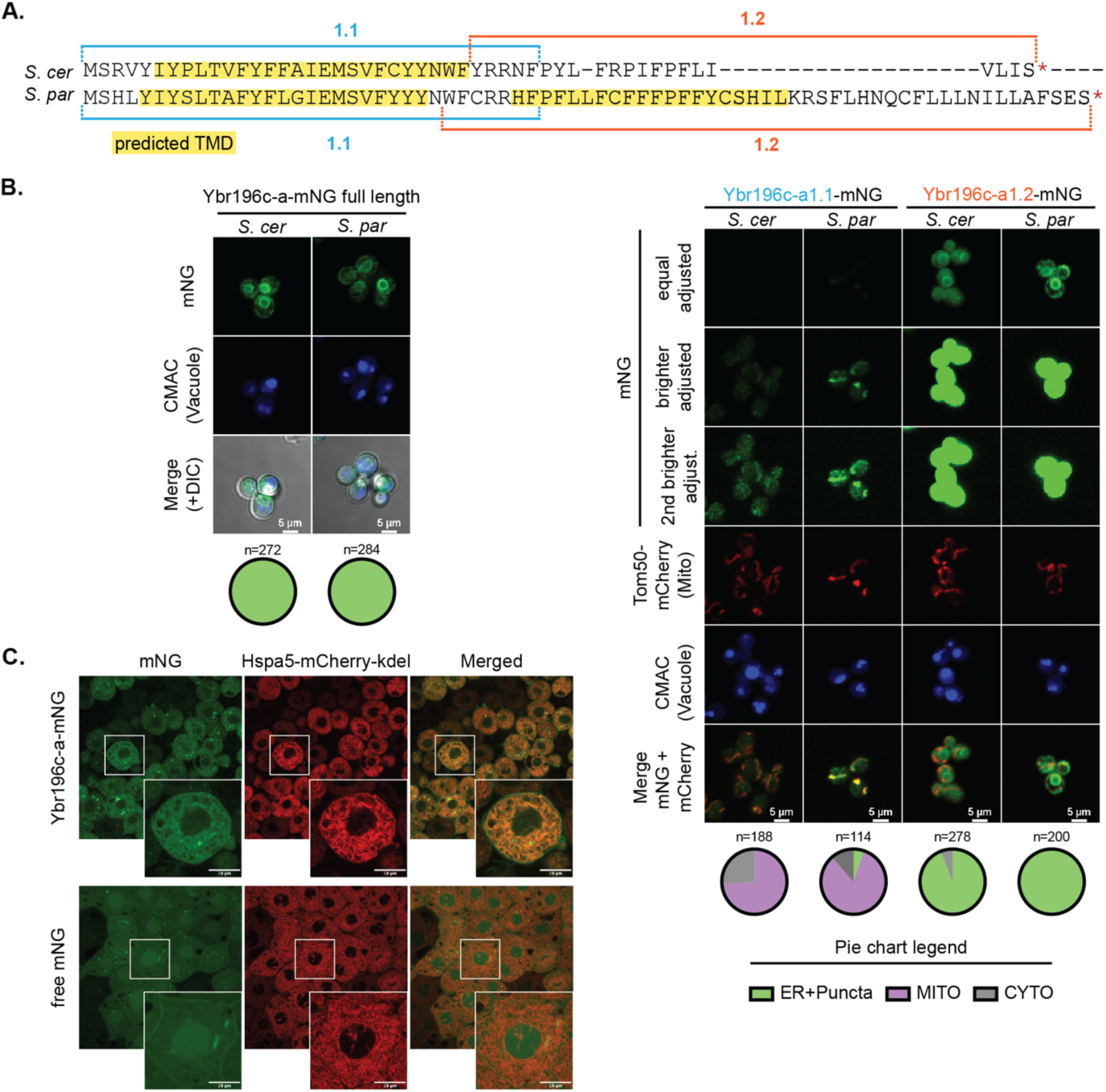
The DNP Ybr196c-a localizes to the ER in diverse eukaryotes. **A.** Sequence alignment shown for Ybr196c-a between *S. cerevisiae* (*S. cer*) and its sister species *S. paradoxus* (*S. par*) as obtained from Vakirlis et al 2020 (12). TMD predictions made with Phobius (18) are shown in yellow. Brackets indicate the sequences used for each subclone, 1.1 and 1.2 in either species. Full-length and partial sequences for both species were fused C-terminally to mNG and genomically integrated into *S. cerevisiae* under the control of an inducible promoter. **B.** The C-terminal region of Ybr196c-a and its orthologous sequence in *S. paradoxus* drive localization to the ER. Fluorescent micrographs are shown of the full-length (left) and two partial sequences (right) of the Ybr196c-a proteins from both species. An mCherry-tagged Tom50 was genomically integrated into these strains to allow visualization of the mitochondria and cells were stained with CMAC-blue to mark the vacuoles. The distributions of localizations in single cells (n) observed for each construct are represented in the pie-chart graphs to the right of each image. For simplicity, in partial sequences (right) the merge shows the co-localization between the mNG and the mCherry signal only. **C.** Representative micrographs from fluorescence microscopy of Ybr196c-a-mNG localization (top) in developing zebrafish embryos. Hsp4 tagged mCherry was used to visual the ER. Controls embryos were injected with mRNA containing only the free mNG (bottom). Insets are shown to visualize colocalization of Ybr196c-a-mNG or free mNG with Hsp4 tagged mCherry.

We posited that if the pathways needed to localize Ybr196c-a were conserved, expression of this DNP in other organisms would result in a similar ER localization. To test this hypothesis, we examined Ybr196c-a in distant taxa. We injected zebrafish embryos in the 1-cell stage with mRNA for Ybr196c-a fused to mNG, together with the ER-marker Hspa5 fused to mCherry and imaged them at the early gastrulation stage (**Figure 3C**). Ybr196c-a co-localized with the ER-marker Hspa5 (**Figure 3C**). These results provide support for the proposition that, while the ER-localized DNPs we investigate here are not themselves conserved, the systems and rules that govern their targeting may be conserved and generally consistent across evolutionary time.

### ER targeting of DNPs depends on conserved post-translational machinery

Our model ER-localized DNP Ybr196c-a might reach the ER via a canonical co- or post-translational targeting mechanisms. We evaluated whether its localization was altered upon perturbing three highly-conserved eukaryotic ER targeting pathways (SRP-dependent, GET, and SND) (25). To this aim, we engineered relevant gene deletions into strains expressing a chromosomally integrated Ybr196c-a-mNG fusion under the control of an inducible promoter, as in **Fig 3B**. Upon disruption of the core GET or SND machinery (Get1-3 or Snd2-3, respectively), fluorescence signal at the ER was reduced and an elevated number of cytosolic puncta was observed (**Fig 4A-C**, **Fig S5A**). This punctate localization pattern is expected for protein clients of these post-translational insertion pathways, as disruption of these targeting mechanisms results in cytosolic aggregation of client proteins (30, 31). In contrast, disruption of GET pathway accessory proteins (*e.g.,* Get4, Get5 and Sgt2) or the co-translational SRP-dependent targeting components (*e.g.,* Sbh1, Sec71, and Sec72) did not alter Ybr196c-a localization (**Fig S5A**). Ybr196c-a’s reliance on operational GET and SND systems for its ER targeting was more apparent with the N-terminal than the C-terminal mNG-tagged forms, as might be expected for a protein that can insert with mixed topology but prefers C-tail anchored protein insertion (**Fig 4A-C**, **Fig S5A**). Perturbing GET or SND components had no impact on free mNG localization, ruling out effects of the fluorescent tag on Ybr196c-a’s targeting (**Fig S5B**). Together, these findings demonstrate that Ybr196c-a depends on functional GET or SND machinery to access the ER.

**Figure 4.**
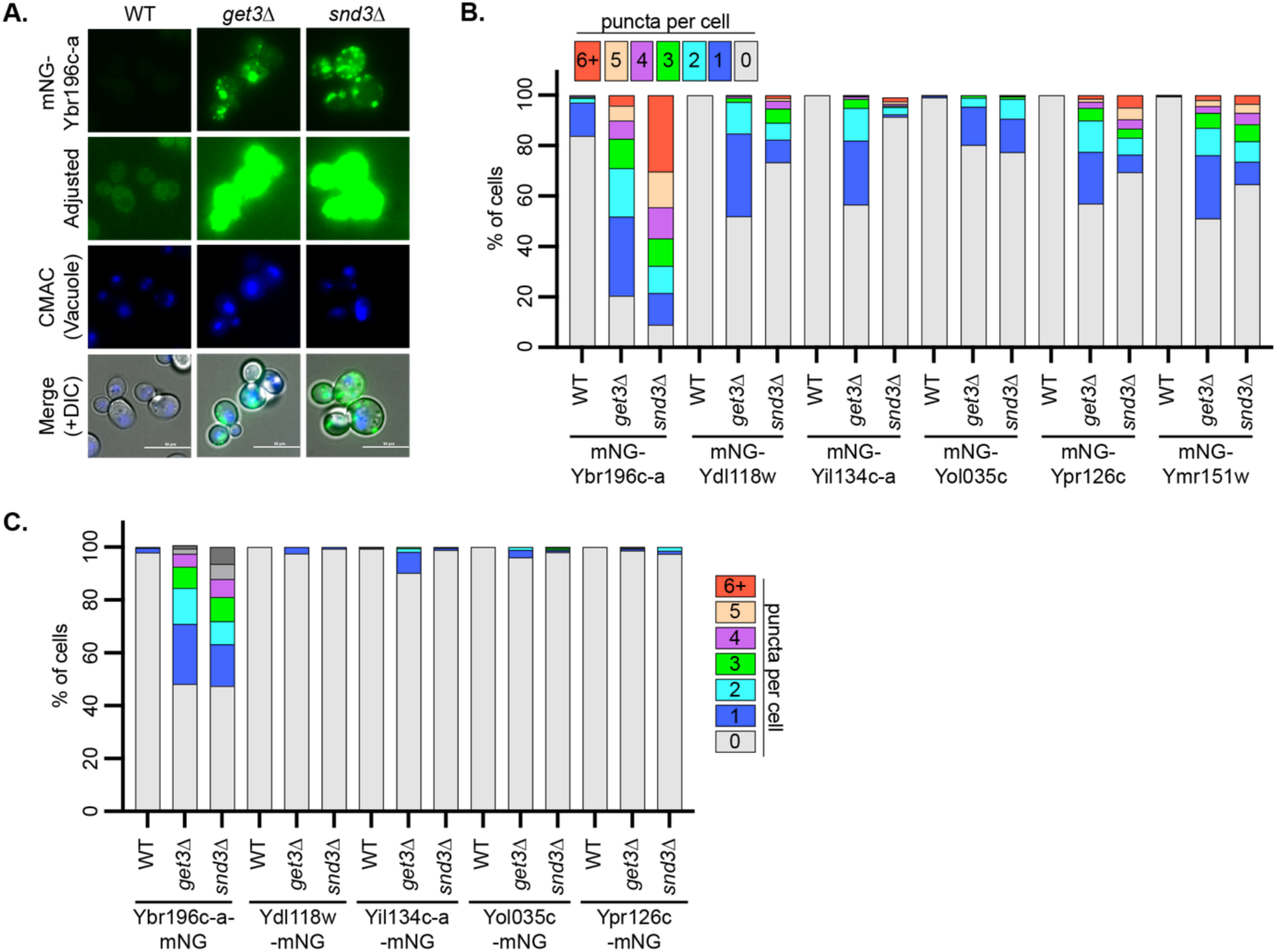
The GET and SND posttranslational targeting pathways are needed for DNPs to access the ER. **A.** Fluorescence micrographs of mNG-Ybr196c-a in cells with the indicated deletions of GET or SND components (Get3 or Snd3, respectively) are shown. **B and C**. Increasing numbers of puncta per cell, and concomitant loss of clear ER localization, are associated with disruption of the ER-targeting pathways needed for DNP targeting. The percentage of cells (n=305-881) with the number of puncta per cell (based on images shown in panel A and Figure S5C-D), are indicated as a stacked distribution.

To determine if these findings generalize to the other ER-localized DNPs, we examined their localization in the absence of Get3 and Snd3 using chromosomally integrated C- or N-terminal mNG-fusions under the control of an inducible promoter. We found that, like Ybr196c-a, all the ER-localized DNPs were dependent on the GET or SND core machinery for ER-targeting (**Fig 4B-C** and **Fig S5C-D**). This effect was more pronounced when the proteins were N-terminally tagged and their C-termini were accessible (**Fig 4B-C**). We hypothesize that this asymmetry is caused by the C-terminal mNG tags masking the TMDs needed for efficient post-translational ER insertion. Indeed, a similar effect as has been reported for other proteins (32) and linked to a favored topology of insertion where the C terminal end more efficiently translocates through the membrane (28). Together these results reveal that all ER-localized DNPs with increased growth overexpression phenotypes depend on the core components of the GET and SND post-translational targeting pathways to reach the ER. Thus, while these DNPs are evolutionarily novel, the GET and SND pathways are conserved across eukarya possibly explaining the conserved ER localization for Ybr196c-a in yeast and zebrafish (33).

### Conserved ERAD pathways regulate DNP stability at the ER

Having established that DNPs depend on conserved pathways for ER localization, we next assessed conserved pathways that might regulate their homeostasis. We first asked whether the abundance of our model ER-localized DNP, Ybr196c-a, depends on the two major ER-resident, E3-<u>ub</u>iquitin (Ub) ligases involved in ERAD, Doa10 and Hrd1 (34). Immunoblotting for Ybr196c-a tagged either N- or C-terminally with mNG revealed a large increase in Ybr196c-a levels upon *DOA10* deletion (**Fig 5A-B**). This increase was not driven by the tag: free mNG was not stabilized by the loss of *DOA10* (**Fig S6A-B**), and a similar increase upon deletion of *DOA10* was observed when Ybr196c-a was fused to an HA tag (**Fig S6C-D**). We then examined the protein turnover of Ybr196c-a by blocking translation with the addition of cycloheximide (CHX) (35) and monitoring protein abundance over time. We performed these CHX-chase assays using: 1) C-terminal mNG-tagged Ybr196c-a (**Fig 5C-D**), 2) free mNG (**Fig S6E-F**), 3) C-terminal HA-tagged Ybr196c-a (**Fig S6G-H**), and 4) N-terminal mNG-tagged Ybr196c-a (**Fig S6I-J**). From these assays, we confirmed that all versions of Ybr196c-a (N- or C-terminally-tagged with either mNG or HA) were stabilized by loss of Doa10. Doa10 ubiquitination of ER-resident proteins often results in their retrotranslocation from the ER and subsequent proteasomal degradation in the cytoplasm (36). Accordingly, we found that Ybr196c-a was more stable in the presence of a proteasome inhibitor than in vehicle control-treated cells (**Fig 5E-F**), unlike free mNG (**Fig S6K**) and irrespective of the positioning of the mNG tag (**Fig S6L**). Together, these experiments demonstrate that the protein stability of our model ER-localized DNP depends on the ER-resident Ub ligase Doa10 and proteasome-dependent degradation.

**Figure 5.**
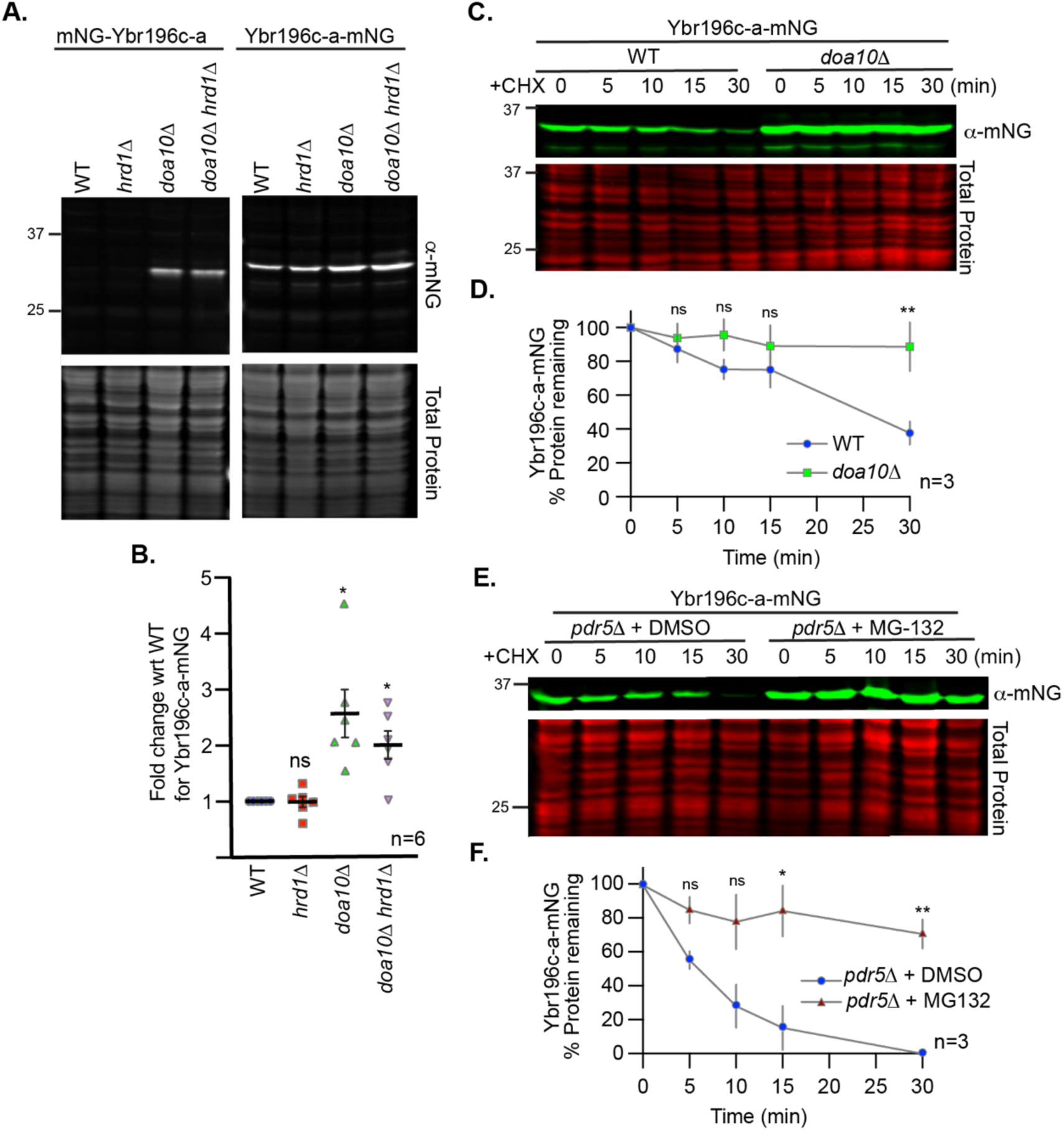
Ybr196c-a’s stability is dependent upon Doa10 and the proteasome. **A.** A representative immunoblot of protein extracts made for N- or C-terminally mNG-tagged Ybr196c-a expressed in the indicated cells. **B.** Quantification of the relative abundance of Ybr196c-a-mNG from six biological replicates of immunoblots, as in **A,** for the cells indicated. A Student’s t-test comparing the WT to the Ub ligase deletions assessed statistical differences. (ns: not significant; *: p-value < 0.05). **C** and **E.** Representative immunoblot of protein extracts expressing Ybr196c-a-mNG from cells post cycloheximide (CHX) (in panel C), or CHX plus MG-132 or DMSO (in panel E) addition for the times indicated (in minutes). When the proteasomal inhibitor MG-132 was added, *pdr5*Δ cells were used to improve retention of MG-132. **D** and **F.** Quantification of the percent of Ybr196c-a-mNG remaining post-CHX addition in the indicated backgrounds. The dots represent the mean of the biological replicates (n), and the error bars show the standard error of the mean. A Student’s t-test comparing WT to the *doa10*Δ cells (in panel D) and DMSO- to the MG-132-treated cells (in panel F) at each time point was used to assess statistical differences (ns: not significant; *: p-value < 0.05; **: p-value <0.005; ***: p-value <0.0005). Similar analysis to those presented in **C** and **D** for an mNG control and the N-terminally mNG-tagged Ybr196c-a are presented in Figures S6E-F, S6I-J, respectively. Similar analysis to those presented in **E** for an mNG control and the N-terminally mNG-tagged Ybr196c-a are presented in Figures S6K and S6L, respectively. For all immunoblots, REVERT total protein stain serves as the loading control.

To confirm our findings and determine if they generalize to the other ER-localized DNPs, we examined their abundance using microscopy in the absence of Doa10 and/or Hrd1 using chromosomally integrated C- and N-terminal mNG fusions under the control of an inducible promoter. Using this approach, we confirmed that Ybr196c-a abundance and ER localization increases in the absence of Doa10, irrespective of tag location (**Fig 6A-B**). Quantification of whole-cell fluorescence from these images revealed Ybr196c-a’s abundance also significantly increased upon loss of Hrd1, but less so than in the Doa10 deletion cells (**Fig 6C**). Like Ybr196c-a, we found that all the ER-localized DNPs increased in abundance and ER localization upon loss of at least one of the Ub ligases; immunoblotting revealed distinct bands (**Fig S7A-E**), and microscopy revealed increased whole-cell fluorescence and ER localization (**Fig S8**). Based on immunoblotting and microscopy, protein stabilization was more pronounced for N-terminally tagged DNPs, which would make their C-termini accessible (**Fig 6D**, **Fig S7-S8**). While a larger fraction of cells with distinguishable ER localization was observed for all C-terminally tagged DNPs upon Ub ligase deletion, whole-cell fluorescence increased for only three of these proteins (**Fig 6D**, **Fig S8**). Collectively these results demonstrate that the homeostasis of all ER-localized DNPs tested depends on the major ER-resident Ub ligases, which are widely conserved in eukarya (37, 38).

**Figure 6.**
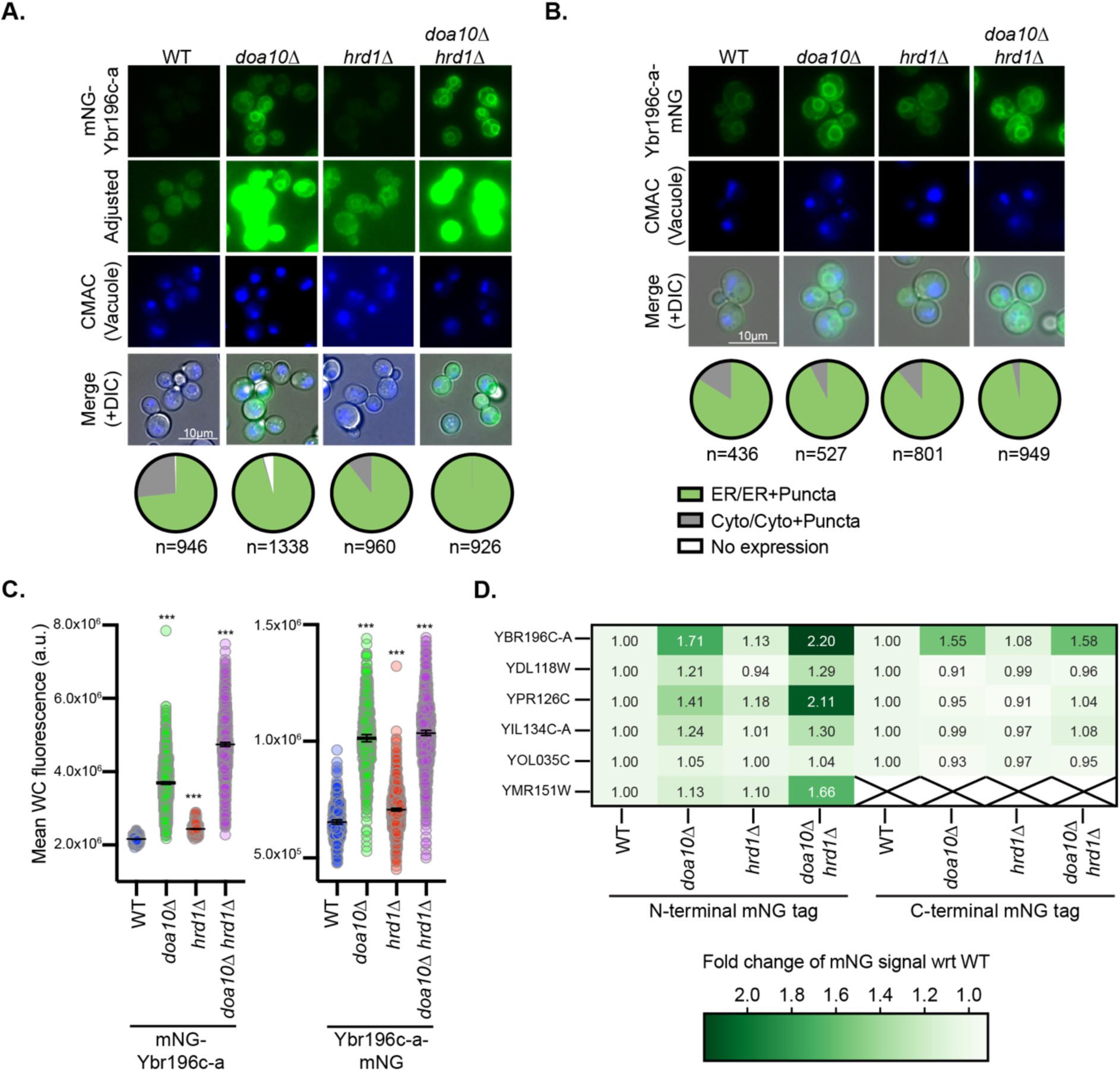
ER-localized DNPs’ abundance depends on Hrd1 and Doa10. **A-B.** Representative fluorescence microscopy of **A**, N-terminally- or **B**, C-terminally-mNG tagged Ybr196c-a. The distribution of fluorescence observed across individual cells (n) is indicated in the pie chart below each image panel. Cells are stained with CMAC blue to mark vacuoles and imaged via DIC, which is shown in the merge. Analogous imaging data for all ER-localized DNPs is provided in Figures S8. **C.** Quantification of the whole cell fluorescence intensities for the cells (n=180-270) shown in panels **A** and **B** is shown. A one-way ANOVA with Dunnett’s multiple comparisons test was performed relative to the fluorescence in WT cells to identify statistically significant changes (p-value < 0.0005 = ***). **D.** A heatmap indicating the fold change in the mNG fluorescent signal for ER-localized DNPs in cells lacking the indicated Ub ligase relative to the WT controls (based on data in C, Figs S8C, S8F, S8H, S8K, S8N).

### Vacuolar protein sorting machinery contributes to DNP homeostasis

Finally, we explored the possibility that the homeostasis of ER-localized DNPs might also be regulated through proteasome-independent mechanisms in the vacuole, the yeast equivalent of the lysosome (39). In the absence of the master vacuolar protease Pep4, fluorescently-tagged proteins targeted to this organelle accumulate vacuolar fluorescence because the fluorophore is not degraded efficiently (40–43). Fluorescence microscopy revealed that all ER-localized DNPs accumulate in the vacuole when Pep4 is deleted, suggesting they are all able to transit the secretory pathway to the vacuole (**Fig 7A-B** and **Fig S9A-G**). The two known transit routes from the ER to the vacuole are via ER-phagy or vesicle-mediated trafficking (44). In the case of vesicular trafficking, disruption of vacuolar protein sorting via loss of Vps4, an ESCRT-III (endosomal-sorting complex required for transport) component (45), will prevent sorting of membrane proteins into multi-vesicular bodies and block their delivery into the vacuole lumen. If ER-localized, DNPs rely on vesicle-mediated trafficking to the vacuole, then loss of Vps4 will block their accumulation in the vacuolar lumen, instead leading to their accumulation in endosomes adjacent to the vacuole (46). Consistently, deletion of Vps4 in cells lacking Pep4 prevented accumulation of mNG signal in the vacuole and led to the formation of large fluorescent puncta adjacent to the vacuole (**Fig 7C-D**, **Fig S9H-L**). These findings imply that ER-localized DNPs traffic through the secretory and/or endocytic pathways beyond the ER. Our results demonstrate that the homeostasis of ER-localized DNPs is not only dependent on the ERAD pathway and the proteasome, but also on the vacuolar protein sorting machinery (**Fig 7E**), which predates the origin of eukaryotes (47).

**Figure 7.**
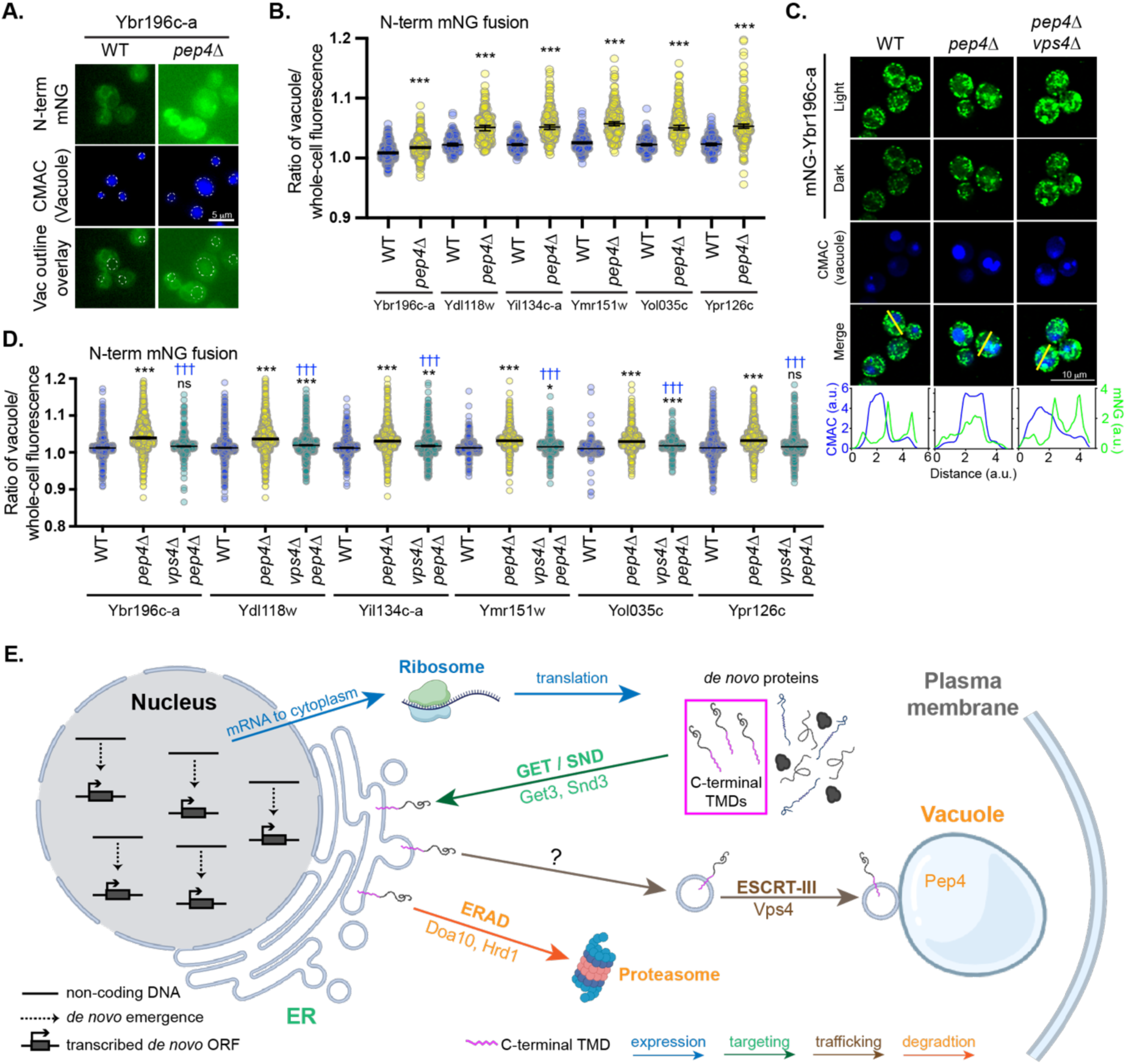
ER-localized DNPs traffic beyond the ER. **A and C.** Fluorescence micrographs of cells expressing N-terminal mNG-tagged Ybr196c-a in WT cells or those lacking *PEP4* and/or *VPS4* are shown. Vacuoles are stained with CMAC-blue. **A**. Vacuoles are indicated on the green channel images using a white dashed line. **C.** Plots indicate the fluorescence intensity from the blue and green channels along a line scan through the vacuole (indicated in yellow on merge). Overlapping maximum peaks of blue and green lines indicate mNG presence in the vacuolar lumen. Similar results and analysis are shown for all other N-terminal mNG-tagged DNPs in Figures S9H-L. **B** and **D.** Scatterplots of the ratio of the vacuolar fluorescence over the whole cell fluorescence for the cells imaged in panel A and Figure S9A are shown in **B** (num of cells=193-924) and those imaged in C and Figures S9H-L are shown in panel **D** (num of cells=206-342). A Mann-Whitney two-sided test assessed statistical difference between the WT and *pep4*Δ (denoted by asterisks) in panel **B** and a one-way ANOVA test with Dunnett’s multiple comparison correction between WT and *pep4*Δ (denoted by asterisks) and *vps4*Δ *pep4*Δ (denoted by blue daggers) in panel **D** cell populations (ns: not significant, p < 0.05 = *, p < 0.0005 = ***, p < 0.0005 = †††). The median fluorescence intensity and 95% confidence interval are represented by horizontal lines. Similar analyses to those shown in panel B for C-terminal mNG-tagged DNPs is provided in Figure S9C. **E.** Model of ER-localized DNP cellular integration. *De novo* originated ORFs arising from non-coding DNA are transcribed and translated into DNPs. ER-localized DNPs all share a common C-terminal TMD (outlined in pink) and depend on the post-translational targeting pathways GET and SND to localize to the ER. Once at the ER, DNPs are either degraded in the proteasome via ERAD, or traffic out of the ER using an unknown mechanism (indicated by a question mark) to eventually arrive at the vacuole via an ESCRT-III-mediated pathway where they are degraded by vacuolar proteases. Specific components of each major pathway found to be involved in DNP targeting, trafficking, and degradation listed below corresponding arrows.

## Discussion

In this study, we report the first experimental investigation of the cellular processing of *de novo* proteins (DNPs). Of the four hypotheses we considered for DNP processing, we find no evidence that DNPs employ noncanonical pathways, operate without assistance from other proteins, or utilize the full set of canonical pathways. However, these results do not rule out these alternative hypotheses, as our study examines only a small subset of recently evolved, growth-enhancing DNPs. Other DNPs may employ processing mechanisms not captured here. Rather, our results demonstrate that a specific subset of ancient targeting, trafficking, and degradation pathways control the homeostasis of yeast DNPs with growth-promoting overexpression phenotypes. Specifically, the recently emerged DNPs we investigated rely on the GET and SND post-translational membrane insertion pathways to localize to the ER and on the ERAD pathways for homeostasis, both of which are conserved across eukarya (33, 37, 38). Similarly, the ESCRT-mediated mechanism that enables trafficking of these DNPs to the vacuole predates the last universal eukaryotic ancestor (47). Our discovery that DNPs are processed by cellular pathways with deep conservation indicates that these pathways can be used by DNPs without an extensive period of co-evolutionary adaptation. The evolutionary conservation of the pathways targeting ER-localized DNPs is further supported by the observation that our model ER-localized DNP also localizes to the ER in zebrafish embryos. Thus, the same pathways may facilitate the integration of DNPs into pre-existing cellular systems across much of the tree of life (2).

Notably, the GET and SND pathways used by all ER-localized DNPs investigated in this study are not the primary mechanism by which most ER-localized ancient proteins are targeted to the ER, with a large majority instead using co-translational mechanisms such as SRP (21). Recently emerged DNPs may not make use of the full range of targeting strategies employed by older proteins, but instead disproportionately depend on select pathways, like GET and SND, that may be easier to access in the absence of long-term evolutionary adaptation. These select pathways, in turn, will have outsized influence on the early evolution of *de novo* genes. The difference in distribution of targeting mechanisms between young DNPs vs. older proteins suggests model whereby young DNPs would use post-translational mechanisms for ER-localization, but would evolve over time to use co-translational mechanisms (21).

We and others have hypothesized that membranes may offer a privileged entry point for DNPs to gain access to the cellular environment (12, 48–51). Several lines of evidence support this idea. For instance, a disproportionate fraction of *de novo* open-reading frames is transcriptionally co-expressed with membrane-associated genes (52). In humans, newly identified microproteins—many of which have evolved *de novo* (53)—exhibit a high prevalence of predicted TMDs (51). In arctic fish, a DNP exerts life-saving anti-freeze activity thanks to its ability to traffic through secretory pathway membranes (54). In yeast, recently emerged DNPs that promote growth when overexpressed under nutrient stress are enriched in predicted TMDs (12). In line with these observations, we find here that most of these growth-promoting DNPs localize to membrane-bound compartments, and particularly the ER. We find that DNPs associated with increased colony size are four times as likely as conserved proteins to localize to the ER. Moreover, ER-localized DNPs are more likely to improve growth when overexpressed than other DNPs. Though we cannot be confident that every protein that is able to improve growth when overexpressed is beneficial, the strong association between ER localization and growth indicates that ER localization is phenotypically consequential and may be particularly likely to mediate adaptive phenotypes. Conserved human ER-localized microproteins often regulate larger protein complexes through protein interactions (55), and so it is plausible that novel ER proteins may function through similar mechanisms.

How do DNPs acquire the signals necessary to target to the ER? Targeting to the ER by the GET and SND pathways is somewhat promiscuous, relying on the recognition of crude molecular signatures (TMDs) (56, 57) that are common in non-coding DNA (12, 58, 59) and accessible without extensive prior adaptation. Generally, non-genic sequences enriched in thymine such as poly dA/dT tracks tend to encode hydrophobic amino acids and have a high propensity to form TMDs (12, 58, 59). It is plausible that DNPs regularly emerge with biochemical properties that support their membrane targeting. For example, Ybr196c-a emerged from a thymine-rich promoter sequence that already possessed the capacity to form a C-terminal TMD and the efficiency of its ER targeting may have increased over time (12). Future studies will be needed to determine the precise evolutionary mechanism by which DNPs acquired their shared capability to access the ER, and whether ER-targeting signals were already present upon *de novo* emergence, under the action of natural selection, or a combination of both (60). We find all ER-localized DNPs are subject to degradation by ERAD and vacuolar proteases. The significance of this finding warrants careful interpretation in the context of *de novo* emergence. High rates of degradation could be construed as a barrier to the emergence of *de novo* genes. Alternatively, regulated degradation may facilitate *de novo* gene birth by mitigating the potentially negative effects of high levels of *de novo* protein expression, while maintaining some molecules to carry out biological activities.

Furthermore, an evolutionary model where the ability to co-opt degradation pathways represents a selective advantage for DNPs would align mechanistically with our discovery that DNPs tend to localize at the ER, an ancient organelle well-equipped for stringent protein quality control of nascent transmembrane proteins. The ER membranes may serve as a “safe harbor” (48) for *de novo* TMD proteins, facilitating their evolutionary persistence by offering a prospective adaptive niche while ensuring that excess proteins are efficiently cleared to mitigate toxicity and reduce purifying selection. Our data show that DNP’s hydrophobic C-terminal TMDs are involved in their ER targeting. Recent screens and computational analyses suggested that hydrophobic C-termini can also act as signals for proteasomal degradation and influence the evolution of human noncanonical translated sequences (49, 50, 61). Thus, the same molecular signal may mediate clearance from the cytoplasm through both degradation and ER localization.

Overall, our findings reveal that young DNPs can display consistent modes of regulation despite having unique primary sequences that are new to nature. DNPs that localize to the ER contain similar molecular signatures, targeting mechanisms, modes of degradation, and phenotypic outcomes, pointing to a common, favored route of integration in the cell. By channeling proteins through specific pathways, cells may bias the molecular features—and thus the functional capacities—of proteins that successfully emerge *de novo*. As a result, the functional repertoire of young genes may initially represent a biased subset of the molecular activities conserved proteins can perform. Our study paves the way toward predictive models linking molecular signatures, cellular integration, and phenotypic impacts of DNPs across species and environmental contexts.

## Materials and Methods

### Plasmids and DNA manipulations

#### Gateway entry clones

To create a collection of entry clones for cloning with the Gateway System (Thermo Fisher, Waltham, MA), primers were designed containing the attB1 and attB2 sequences to amplify each ORF with a stop codon for the N-terminal tagging and without, for the C-terminal tagging. The primers were synthesized by Integrated DNA Technologies (IDT, Coralville, IA) and used to amplify each ORF using Q5 High-Fidelity 2x Master Mix (M0492L, New England BioLabs, Ipswich, MA). Genomic DNA from the strain FY4 was extracted with a Yeast DNA Extraction kit (78870, Thermo Fisher) and used as a template for the PCR. The PCR conditions were as follows: 98°C for 30 s, followed by 25 cycles of 98°C for 10 s, 55°C for 15 s and 72°C for 30 s and a final elongation step of 2 min at 72°C. The PCR products were purified with NucleoFast 96 PCR Plates (743100.10, Macherey Nagel, Allentown, PA) and quantified on a Nanodrop (Thermo Fisher) prior to use for recombination with the donor plasmid pDONR223, using BP Clonase II Enzyme mix (11789100, ThermoFisher). The recombination reactions were used to transform DH5a competent cells (C2987U, New England BioLabs) and positive clones were selected in LB media(62) supplemented with 50 µg/ml of spectinomycin (158993, MP Biomedicals, Santa Ana, CA). The NucleoSpin 96 Plasmid kit (740625.4, Macherey Nagel) was used to extract the plasmids and DNA quantification was done using the SpectraMax M4 plate reader (Molecular Devices, San Jose, CA).

#### Destination plasmids

Two sets of destination plasmids were used in this study. The first set from the Yeast Gateway kit (1000000011, Addgene, Watertown, MA(63)) includes the plasmids pAG415-GAL-ccdB-EGFP and pAG415-GAL-EGFP-ccdB. The second set was built to swap the GAL promoter with the Z_3_EV promoter and the ccdB-EGFP was replaced with ccdB-mNG-Tadh1 and a hygromycin cassette (for the C terminally tag) or mNG-ccdB-Tadh1and a hygromycin cassette (for the N terminally tag), creating the destination plasmids pARC0031 and pARC0152.

#### Expression plasmids

Expression plasmids were made by LR recombination between the entry clones and the destination plasmids described above, using the LR Clonase II enzyme mix (11791020, Thermo Fisher). The recombination reactions were used to transform DH5a competent cells (C2987U, New England BioLabs) and positive clones were selected in LB media supplemented with 100 mg/ml of ampicillin (J60977, Alfa Aesar, Haverfill, MA). The NucleoSpin 96 Plasmid kit (740625.4, Macherey Nagel) was used to extract the plasmids and DNA quantification was done using the SpectraMax M4 plate reader (Molecular Devices). We attempted to create expression plasmids for all 28 DNPs from Vakirlis et al (12), but only succeeded in constructing a final plasmid containing a DNP-EGFP fusion for 26 of the DNPs (YGR296W and YOR011W-A failed). Plasmids used in this work are described in **Supplemental Table 2**.

#### Yeast strains and growth conditions

The yeast strains used are described in **Supplemental Table 3** and are all derived from S288c genetic backgrounds of *S. cerevisiae*.

Yeast cells were grown in either <u>s</u>ynthetic <u>c</u>omplete medium (SC) lacking the appropriate amino acids for plasmid selection, prepared as described in (64) and using ammonium sulfate as a nitrogen source, or YPD (62) medium where indicated. Liquid medium was filter-sterilized and solid plate medium had 2% agar (w/v) added before autoclaving. When necessary for selection, hygromycin B or G-418 (H75020-1.0 and G64000-5.0, Research Products International, Mt Prospect, IL) was added to the media to a final concentration of 200 µg/ml. Yeast cells were grown at 30°C and where appropriate, 10-20 µM of β-estradiol (E2758-1G, Millipore-Sigma, St. Louis, MO) was added to cultures for 3 hours to induce expression from the *GEV* or *Z*_3_*EV* promoters (15).

#### Yeast transformation

Strains containing plasmids overexpressing fusions with EGFP were transformed in the BY4741 background with or without *ACT1pr*-Z_3_EV-NatMX to be used with β-estradiol or galactose induction, respectively. All other strains were built based on the DBY12394 containing the *ACT1pr*-Z_3_EV-NatMX expression system generously provided by the Noyes lab (15).

The constructs or DNA cassettes amplified from plasmids were integrated in the genome using the lithium acetate (LiAc), polyethyleneglycol (PEG) and salmon sperm DNA transformation protocol (65) with an adaptation for high-throughput scale. The yeast strain was grown in 1ml of YPD media in 96-deep-well plates and used for transformation with the liquid-handler EVO 150 (Tecan Group Ltd, Switzerland). Cells were washed in 750 µl of water, followed by 1 ml of LiAc tris-EDTA (TE) buffer and finally resuspended in 500 µl of the same buffer. Transformations were carried out using 5 µl of ssDNA (2 mg/ml) with 1 µg of purified PCR product or 500 ng of plasmid and 50 µl of cells. 250 µl of PLATE (40% PEG, 0.1M LiAc, 10mM Tris-HCl, 1mM EDTA) was added to each cell mixture and incubated at 30°C for 15 min, followed by 60 min at 42°C. Cells were then pelleted, resuspended in 200 µl of YPD or SC media for a recovery step at 30°C for 3 h, prior to being plated as 3 µl drops on selective medium. Plates were incubated at 30°C for 2-4 days until transformants grew. Agar plates were then used to pin into liquid media with the Singer RoToR (Singer Instruments, Somerset, UK) and then used to make glycerol stocks.

The constructs containing the *Z_3_EVpr* followed by the coding sequences for individual Ds tagged with mNG at the N- or C-terminus were amplified using the expression plasmids as template and integrated at the *HIS3* chromosomal locus. Amplification was carried out using Q5 High-Fidelity 2x Master Mix (M0492L, New England BioLabs) and primers ARC338 (TCTATATTTTTTTATGCCTCGGTAATGATTTTCATTTTTTTTTTTCCACCTAGCGGATGACTCTTTTTTTTTCTTAGCGATTGGCATTATCACATAATGAATTATACATTATATAAAGTAATGTGATTTCTTCGAAGAATATACTAAAAAATGAGCAGGCAAGATAAACGAAGGCAAAGacaaaagctggagctctagta) and ARC339 (AAAGAAAAAGCAAAAAGAAAAAAGGAAAGCGCGCCTCGTTCAGAATGACACGTATAGAATGATGCATTACCTTGTCATCTTCAGTATCATACTGTTCGTATACATACTTACTGACATTCATAGGTATACATATATACACATGTATATATATCGTATGCTGCAGCTTTAAATAATCGGTGTCAgcgaattgggtaccggcc) in a 50 ml reaction. The PCR conditions were as follows: 98°C for 30 s, followed by 25 cycles of 98°C for 10 s, 56°C for 15 s and 72°C for 2 min and a final elongation step of 5 min at 72°C. The PCR products were incubated at 37°C with 1µl of *Dpn*I (R0176S, New England BioLabs) and then purified with NucleoFast 96 PCR Plates (743100.10, Macherey Nagel) and the DNA was quantified on a Nanodrop (Thermo Fisher) prior to being used for yeast transformation. We were not successful in integrating YLL006W with either an N- or C-terminal mNG, nor for YMR151W with a C-terminal mNG, leaving 6 DNPs with mNG tagged on the N terminus and 5 with a C-terminal mNG tag.

Gene deletions were performed with the same transformation protocol and using DNA cassettes (*URA3* or *KANMX*) targeted to the region of interest using primers containing sequences homologous to the genomic locus.

#### Yeast protein extraction and immunoblot analyses

Whole cell extracts of yeast proteins were generated using trichloroacetic acid (TCA) method as described in (66) and modified from (67). In brief, cells were grown in SC medium to mid-exponential log phase at 30°C (A_600_ = 0.6-1.0) and an equal density of cells was harvested by centrifugation. Cell pellets were frozen in liquid nitrogen and stored at -80°C until processing. Cells were lysed using sodium hydroxide, precipitated with 50% TCA, solubilized in SDS/Urea buffer [8 M Urea, 200 mM Tris-HCl (pH 6.8), 0.1 mM EDTA (pH 8.0), 100 mM DTT, 100 mM Tris (not pH adjusted)] and heated to 37°C for 15 min prior to loading on an SDS-PAGE gel. Immunoblotted proteins were detected using mouse anti-green fluorescent protein (GFP) (Santa Cruz Biotechnology, Santa Cruz, CA), mNeonGreen (Cell Signaling Technology, Danvers, MA) or HA antibodies (Roche, Indianapolis, IN). Anti-mouse secondary antibodies conjugated to IRDye-800 or IRDye-680 were used to detect primary GFP or mNG antibodies on an Odyssey CLx infrared imaging system (LI-COR Biosciences, Lincoln, NE). The HRP-conjugated anti-HA antibody was detected on a ChemiDoc (Bio-Rad, Hercules, CA). As a loading and transfer control, membranes were stained with Revert (LI-COR Biosciences) and detected using the Odyssey CLx.

Extracts containing HA-tagged DNPs were loaded on 16.5% Tris-tricine gels and run in the cold using 1X tricine running buffer (Bio-Rad). Proteins were blotted to Immobilon-PSQ (0.2 micron) PVDF membrane using the Criterion system (Bio-Rad). Membranes were stained with REVERT total protein stain (LI-COR Biosciences), followed by blocking with TBST with 3% BSA and overnight incubation at 4°C on a platform rocker with an HRP-conjugated anti-HA antibody in TBST containing 1 % BSA (1:5000 dilution, Roche). Membranes were then washed 3 times with TBST and detected using the SuperSignal West Pico PLUS Chemiluminescent Substrate (Thermo Fisher) on the Bio-Rad Chemidoc XRS+ Imaging System (Bio-Rad).

#### Yeast protein stability assays

The stability of mNG- or HA-tagged YBR196c-a (as an N-terminal or C-terminal fusion) or mNG alone expressed as a chromosomal integration from the *Z_3_EVpr* was assessed by cycloheximide (CHX) chase assay as described in (35). Cells were grown to mid-exponential phase and induced to express the tagged YBR196c-a or mNG using β-estradiol. Cells were next treated with 150 mg/ml CHX (Gold Bio, St. Louis, MO) and equal densities of cells were harvested at the indicated times. Proteins were extracted from cell pellets and subjected to SDS-PAGE and immunoblotting, as described above.

For assays that employed the proteasome inhibitor MG-132 (Thermo Fisher), cells were incubated with 10 µM of MG-132 (stock 10 mM in DMSO) or an equivalent volume of DMSO (vehicle control) for 1 h prior to CHX addition. When CHX was added to block new protein synthesis the t=0 timepoint was harvested to initiate the time course.

#### Zebrafish husbandry and fish lines

Zebrafish (*Danio rerio*) were maintained according to standard laboratory conditions (68). Zebrafish embryos were raised at 28°C in embryo media (250 mg/l Instant Ocean salt in distilled or reverse osmosis-purified water, adjusted to pH 7.0 with NaHCO_3_). All experiments were performed with TLAB wild-type embryos.

#### Zebrafish mRNA synthesis and microinjections

The coding sequence of the yeast DNP YBR196C-A fused to mNG on the C-terminus was cloned into a pCS2+vector with an upstream SP6 promoter and downstream SV40 polyadenylation signal. Briefly, YBR-196C-A-mNG was amplified with PCR primers (Invitrogen, Waltham, MA) and OneTaq polymerase (New England BioLabs) and gel extracted with the GeneJET kit (Thermo Fisher). pCS2+vector was linearized via PCR and YBR-196C-A was cloned into the linearized pCS2+vector using Gibson Assembly cloning (New England BioLabs). The first 16 amino acids of the *D. rerio* Hspa5 and a KDEL ER retention signal were used to stably target mCherry to the ER (69). The DNA sequence of first 16 amino acids of Hspa5 and mCherry-KDEL was synthesized and cloned into the into the pCS2+vector (Twist Biosciences, South San Francisco, CA). All plasmids were transformed into DH5-alpha bacterial competent cells (New England BioLabs) and grown in LB supplemented with 50 mg/ml ampicillin (Thermo Fisher). Single colonies for all plasmids were isolated and confirmed by whole plasmid sequencing (Plasmidsaurus, Louisville, KY). Messenger RNA for YBR196C-A-mNG and Hspa5-mCherry-KDEL was produced by linearizing the template plasmid using NotI transcribing using the mMESSAGE mMACHINE SP6 kit (Invitrogen). mRNA was purified using the Monarch Spin RNA Cleanup Kit (New England BioLabs). All nucleic acid quantifications were carried out with a NanoDrop (Thermo Fisher).

For microinjections, embryos were collected in embryo media and dechorionated using 1 mg/ml Pronase (Protease type XIV from *Streptomyces griseus*, Roche) at one-cell stage. Microinjections were performed on dechorionated one-cell embryos using the Drummond Nanoject III injector. 100 pg of each mRNA as well as phenol red was injected into each embryo (1nL total volume per embryo). Dechorionated embryos were incubated in 1% agarose coated 6-well plates at 28°C until imaging at early gastrulation stages (4-5 hours post fertilization).

### Fluorescence microscopy

#### Yeast imaging

For imaging experiments, cells were grown and induced to express mNG-tagged DNPs or free mNG as indicated above. Fluorescent proteins were localized using: 1) epifluorescence microscopy, 2) confocal microscopy in low-throughput, or 3) confocal microscopy in high-throughput. For epifluorescence microscopy and low-throughput confocal microscopy, cells were stained with 70 µM Cell Tracker Blue CMAC dye (Life Technologies, Carlsbad, CA) and 10 µM trypan blue (Gibco, Dublin, Ireland) and plated onto 35 mm glass bottom microwell dishes that were concanavalin A (MP Biomedicals, Solon, OH) or poly-D-lysine coated (MatTek Corporation, Ashland, MA). For epifluorescence microscopy, cells were imaged using a Nikon Ti2 inverted microscope (Nikon, Chiyoda, Tokyo, Japan) outfitted with an Orca Flask 4.0 cMOS camera (Hammamatsu, Bridgewater, NJ) and a 100x objective (NA 1.49). For low-throughput confocal microscopy, images were captured using a Nikon Ti2 A1R inverted microscope (Nikon) equipped with a 100x objective (NA 1.49) and detected with GaAsP or multi-alkali photomultiplier tube detectors. For high-throughput confocal microscopy, imaging was done as described in Bowman *et al*. (70).

In all cases, image acquisition was controlled using NIS-Elements software (Nikon) and all images within an experiment were captured using identical settings. Images were cropped and adjusted equivalently using NIS-Elements (Nikon), and where added adjusted images are needed to capture the range of fluorescence intensities in an experiment, additional images are presented that are also evenly adjusted and the change in this adjustment is indicated in the figure panel.

#### Zebrafish imaging

Embryos were staged and mounted under a dissecting microscope in 1% low melt agarose in a MatTek dish with a #1.5 glass coverslip. Live imaging was performed on the Nikon A1 confocal inverted microscope (Nikon) at 100X oil objective on GFP and mCherry channels.

### Fluorescence microscopy image analysis and statistical tests

#### Yeast image analysis

To define the localizations represented for both EGFP- and mNG-tagged DNPs, all images were analyzed using Image J software (National Institutes of Health, Bethesda, MD) with the Cell Counter plugin to categorize the localization for every cell in a field. These manually defined localization patterns are summarized in the pie charts presented in **Figures 1B, 3B, 5A-B, S8A-B,D-E,G,I-J, and L-M**.

To quantify the whole cell fluorescence intensity for mNG-tagged DNPs, we used Nikon NIS-Elements .*ai* (Artificial Intelligence) and Nikon General Analysis 3 (GA3) software packages. We trained imaging on a ground-truth set of samples, where cells were segmented using images acquired with the trypan blue-stained cell surfaces or the DIC images. Next, the NIS.*ai* software performed iterative training until a training loss threshold of <0.2 was obtained, which is indicative of a high degree of agreement between the initial ground truth and the output generated by NIS.*ai*. Fields of images were then processed, using the automated segmentation, and the mean fluorescence in the green channel (480 nm) was measured for each cell. Any partial cells at the edges of images were removed. Fluorescence intensities are plotted as scatter plots using Prism (GraphPad Software, San Diego, CA). We performed one-way ANOVA statistical tests with Dunnette’s post hoc correction for multiple comparisons, and or Mann-Whitney tests when only two samples were compared. In all cases, significant p-values from these tests are represented as: * <0.05; ** <0.0005, *** <0.0005; ns >0.05.

#### Biochemical properties, transmembrane domain prediction, and statistics

Amino acid sequences of the DNPs were analyzed for their biochemical properties using Python. A custom script that used the packages Biopython and ‘peptides’ was used to calculate hydrophobicity, percent charged residues, and length in different sequence regions. Transmembrane domain prediction was conducted by accessing the Phobius (https://phobius.sbc.su.se) and TMHMM 2.0 (https://services.healthtech.dtu.dk/services/TMHMM-2.0/) online servers (19). A FASTA file containing all the amino acid sequences was uploaded and both analyses were run using default parameters. All statistical tests were performed in Python using scipy.stats.

#### Description and re-analysis of published datasets

Localization annotations for ancient proteins (shown in **Fig 1C**) were obtained from a published study using the yeast GFP collection (16). The localizations assignments from replicate “WT1” were parsed and proteins were grouped into “ER” or “other” categories. For proteins that exhibited more than one localization, if one was ER, than that protein was counted as “ER”, otherwise it was placed in the “other”.

Fitness measurements for DNPs (shown in **Fig 1D**) were obtained and reanalyzed from (12). For each DNP we counted the number of conditions where colony size grew larger relative to a control, with the minimum and maximum number being 1 and 5, respectively.

## Data Availability

All raw values and statistics for imaging in this study and Supplementary table 1-3 are available for download: https://github.com/CarvunisLab/ER_DNPS_2025

## Code Availability

All custom scripts to run statistical analysis and analyze DNP sequences is available for download: https://github.com/CarvunisLab/ER_DNPS_2025

## Acknowledgments

We thank Dr. Jeff Brodsky and his lab (Univ. of Pittsburgh) for their discussions of this work. We further acknowledge the technical assistance of Dr. Simon Watkins (Director of the Center for Biologic Imaging, Dept of Cell Biology, Univ. of Pittsburgh), Christina Goldbach (Nikon Instruments Inc.) and Dr. Chaowei Shang (Director of Microscopy and Imaging, Dietrich School of Arts and Sciences, Univ. of Pittsburgh) to establish the high-content imaging approaches used in this work. Additionally, we would like to thank all the members of the Carvunis and O’Donnell labs, as well as, the Pittsburgh Area Yeast Meeting group (PAYM) and Molecular Evolution Laboratory Discussion group (MELD), for their feedback and suggestions on various stages of this work. Figures 1A and 7E were created using BioRender.com. This work was supported by funds provided by the National Institute of General Medical Sciences of the National Institutes of Health grant DP2GM137422 awarded to A.-R.C. and the National Science Foundation grant MCB-2144349 awarded to A.-R.C, a subcontract from this grant awarded to A.F.O. and start-up funds provided by the Dept. of Biological Sciences, Univ of Pittsburgh to A.F.O.

## Abbreviations

DNP: *de novo* Proteins
ER: Endoplasmic Reticulum
SND: SRP-iNDependent targeting
SRP: Signal recognition particle
GET: Guided Entry of Tail-anchored proteins
TA: Tail-Anchored
CTE: C-Terminal Extension
CHX: Cycloheximide
EGFP: Enhanced Green Fluorescent Protein
mNG: m-NeonGreen
TMD: Transmembrane Domain
Ub: Ubiquitination
ERAD: ER-associated degradation
ESCRT: Endosomal-sorting complex required for transport

## Supplemental Figures

**Figure S1.**
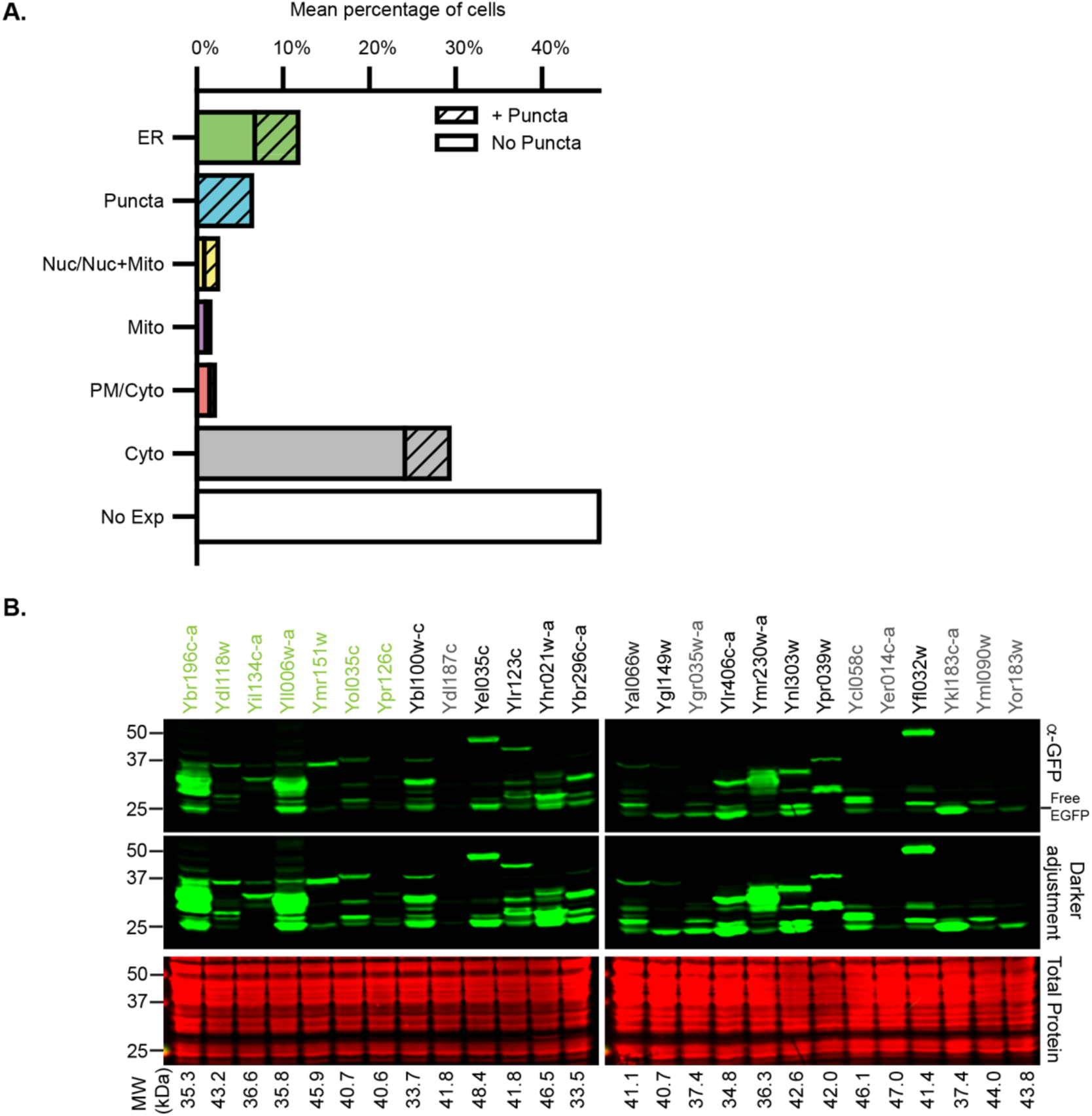
The ER is the predominant membrane localization for DNPs and immunoblotting reveals variable susceptibility to protein processing. (accompanies Figure 1) **A.** The percentages for each localization from Figure 1B (pie charts) are aggregated here to display the mean percentage of individual localizations in the DNP-EGFPs fusions. Membranous compartments on the plot include ER, Nuc (nucleus), Mito (mitochondria), PM (plasma membrane). **B.** Representative immunoblot of protein extracts made for EGFP C-terminally tagged DNPs expressed from plasmids as shown in Figure 1B. Labeled in green text are the ER-localized DNPs, and this designation is based on the microscopy shown in Figure 1B. Expected sizes of the DNP-EGFP fusions indicated below immunoblot. The DNPs in gray are the cases identified as potentially being break down/processed products because the only bands observed are close to the size of free EGFP (∼26.9kDa). REVERT total protein stain serves as the loading control.

**Figure S2.**
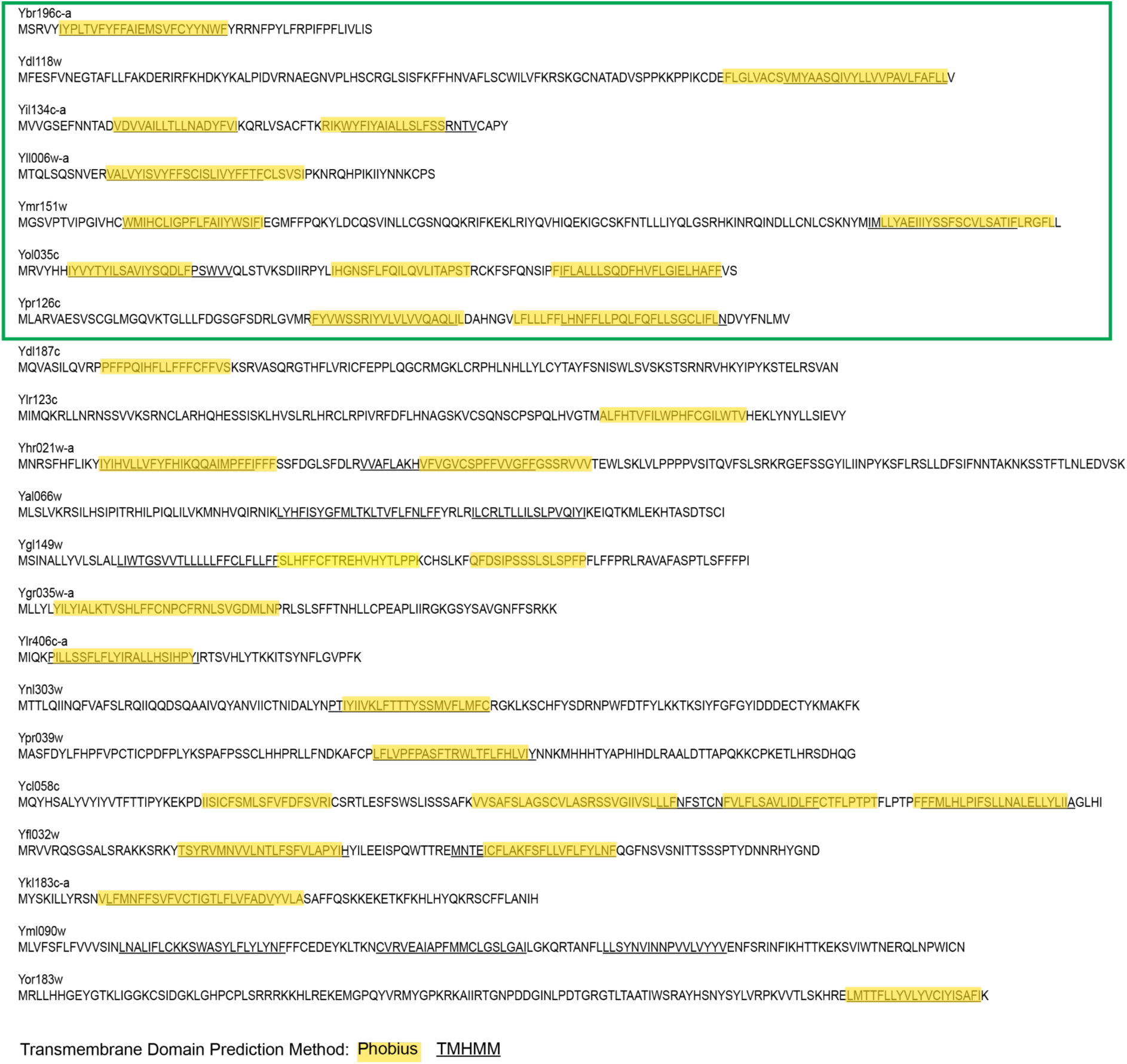
DNPs amino acid sequences and predicted TMDs. (accompanies Figure 2) Amino acid sequences of 21 of the 26 DNPs and those that have at least one predicted TMD, as determined by Phobius (18) or TMHMM (19) are highlighted in yellow or underlined, respectively. The first seven sequences boxed in green correspond to the ER-localized DNPs, which were defined by the microscopy in Figure 1B.

**Figure S3.**
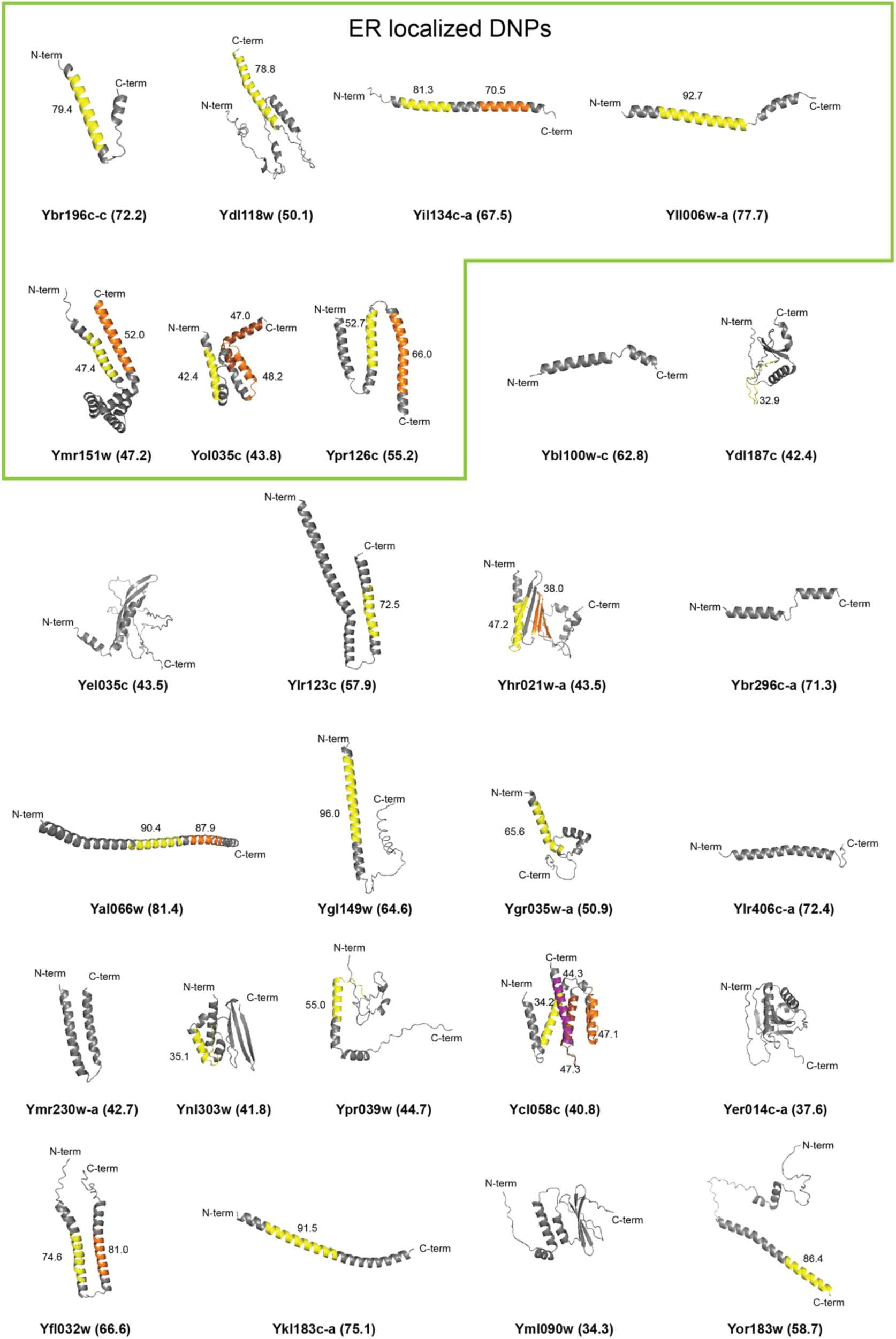
DNPs are rich in alpha helices that correspond with predicted TMDs. (accompanies Figure 2) Alpha-fold 2.0 (20, 71) predicted tertiary structures for the ER-localized (outlined at top) and other DNPs. The N- and C-termini for each DNP are indicated. The confidence of the Alpha-fold 2.0 prediction for each DNP sequence is presented in parenthesis. Note that some Alpha-Fold 2.0 predictions have low confidence due to DNPs’ lack of homology with known proteins (72, 73). TMDs predicted by Phobius (18) are indicated by yellow, orange, and magenta regions and the prediction confidence is presented next to each putative TMD.

**Figure S4.**
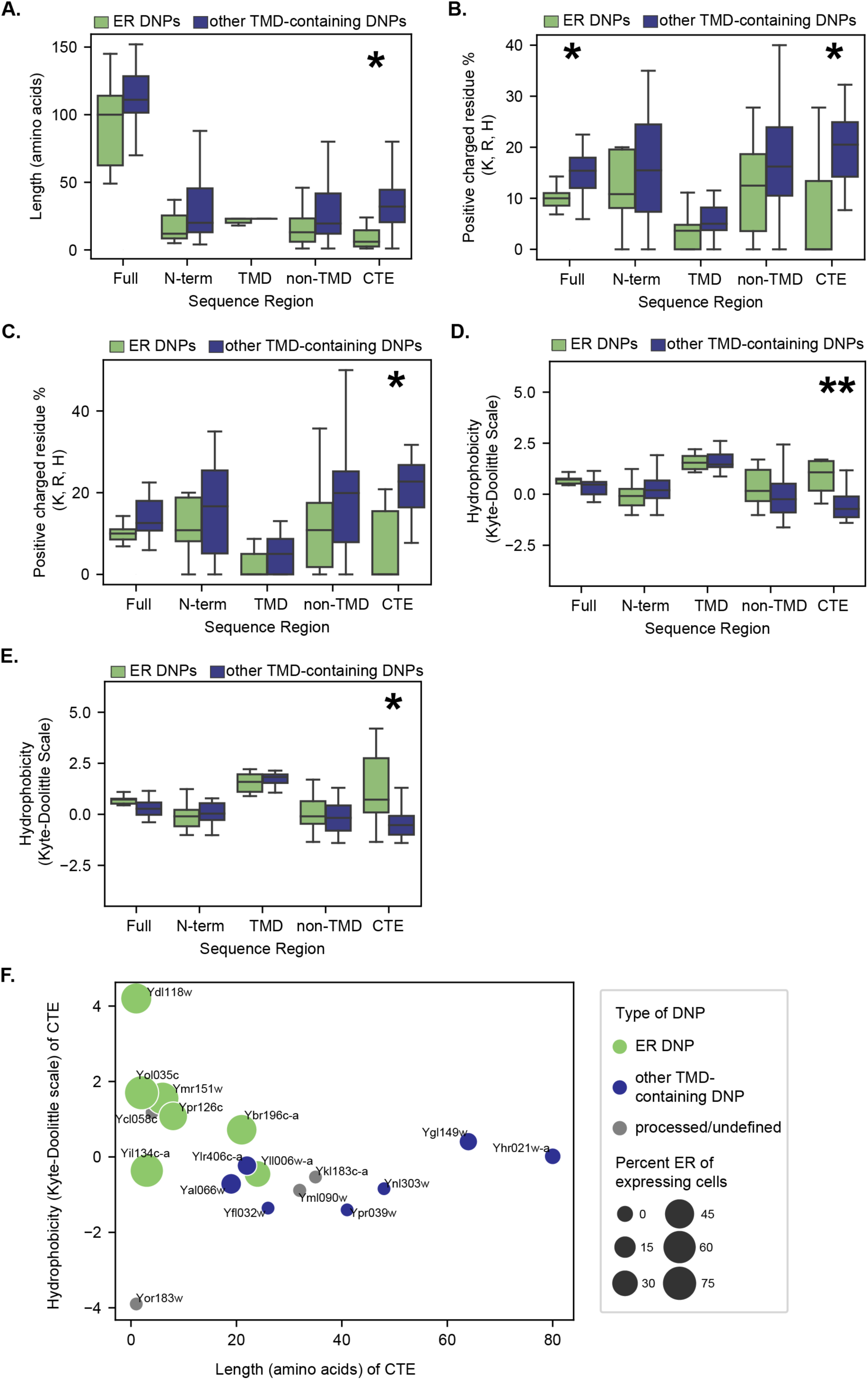
Sequence analysis of TMD-containing DNP protein sequences using Phobius or TMHMM reveals ER-localized DNPs have short, hydrophobic, and less positively charged CTEs. (accompanies Figure 2). **A.** ER-localized DNPs have significantly shorter CTEs than other TMD-containing DNPs. Comparison of the distributions of length in amino acid residues for different sequence regions in ER-localized and other TMD-containing DNPs as predicted by TMHMM (19) are shown. Similar results using Phobius (18) predications are shown in Figure 2B. **B-C.** ER-localized DNPs have less positively charged CTEs than other TMD-containing DNPs. Percent of positively charged residues for sequence regions in both ER-localized and other TMD-containing DNPs as predicted by **B**, Phobius (18) or **C**, TMHMM (19). **D-E.** ER-localized DNPs have significantly more hydrophobic CTEs than other TMD-containing DNPs. Comparison of the distributions of the mean hydrophobicity for different sequence regions in ER-localized and other TMD-containing DNPs as predicted by TMHMM (19) and Phobius (18) are shown in **D** and **E**, respectively. **A-E.** All P values were calculated from Mann-Whitney U test. **: p <0.005, *: p<0.05 Full=Full sequence, N-term=N-terminal region, CTE=C-terminal extension. Full, N-term, and CTE **A, C,** and **D,** n=18 and **B** and **E**, n=19. For TMD **A, C,** and **D**, n=28 and **B** and E, n=30. For non-TMD **A, C,** and **D**, n=46 and **B** and **E**, n=49. **F.** Hydrophobicity versus length in amino acid residues for the CTE of all TMD containing DNPs as predicted by TMHMM (19). Processed/undefined: DNPs that showed some evidence of breakdown in western blots (see Figure S1B). Size of the dots represents percent of expressing cells that had ER localization in Figure 1B. Corresponding systematic gene names from *Saccharomyces* Genome Database are shown next to each dot.

**Figure S5.**
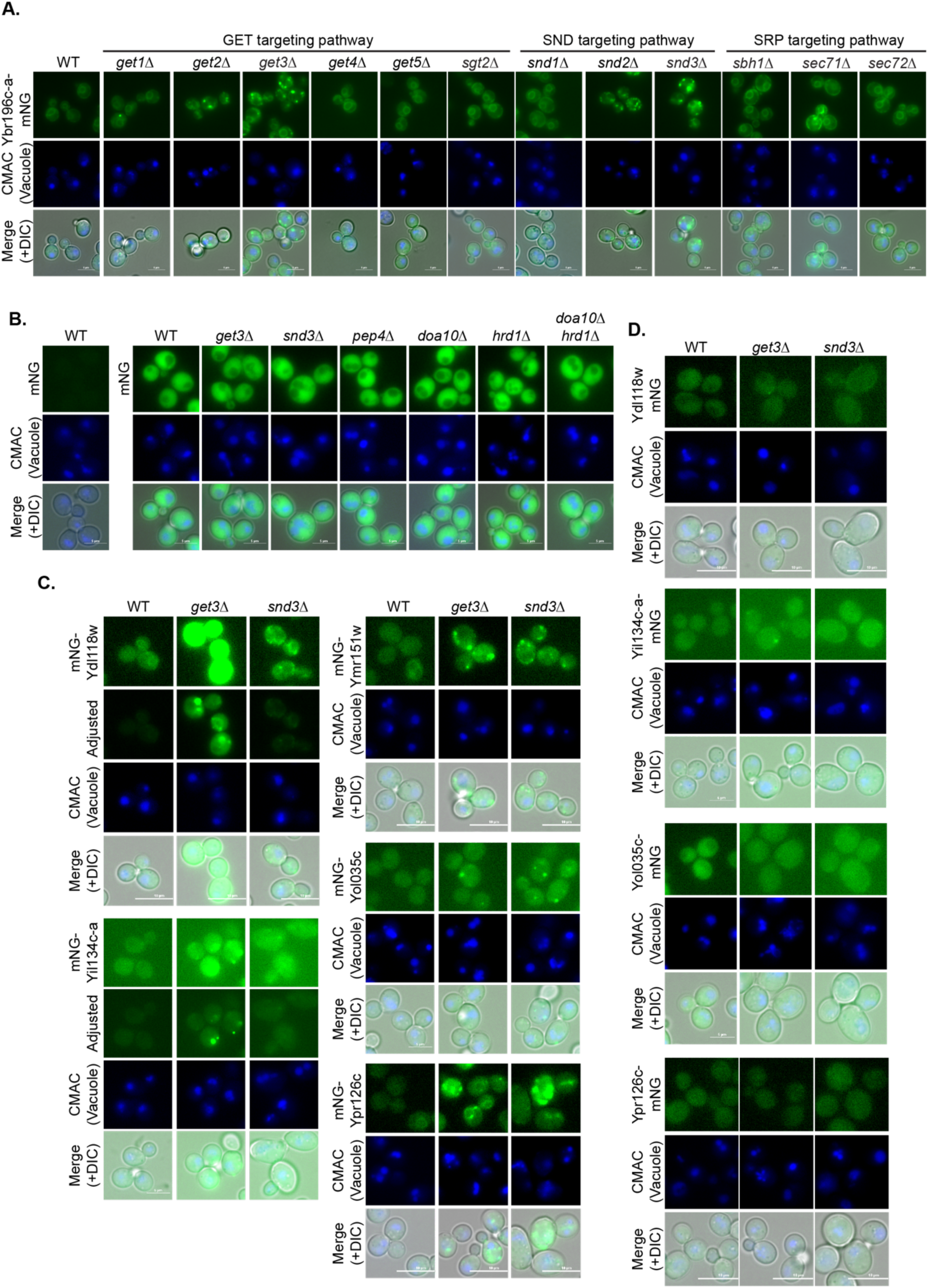
Free mNG and ER-localized DNPs in cells lacking functional GET or SND pathways. (accompanies Figure 4) Fluorescence micrographs of **A.** Ybr196c-a-mNG or **B.** free mNG expressing cells that lack core components of the GET (Get1-3) or SND (Snd1-3) pathways, or GET pathway accessory proteins (*e.g.,* Get4, Get5 and Sgt2) or SRP-dependent targeting components (Sbh1, Sec71, and Sec72) are shown. **C-D.** Fluorescence micrographs of ER-localized DNPs that are N- or C-terminally tagged with mNG, in panels C and D, respectively, are shown. The percentage of cells with the indicated number of puncta per cell are indicated as a stacked distribution in Figure 4B-C, respectively, are based on the images shown in these panels.

**Figure S6.**
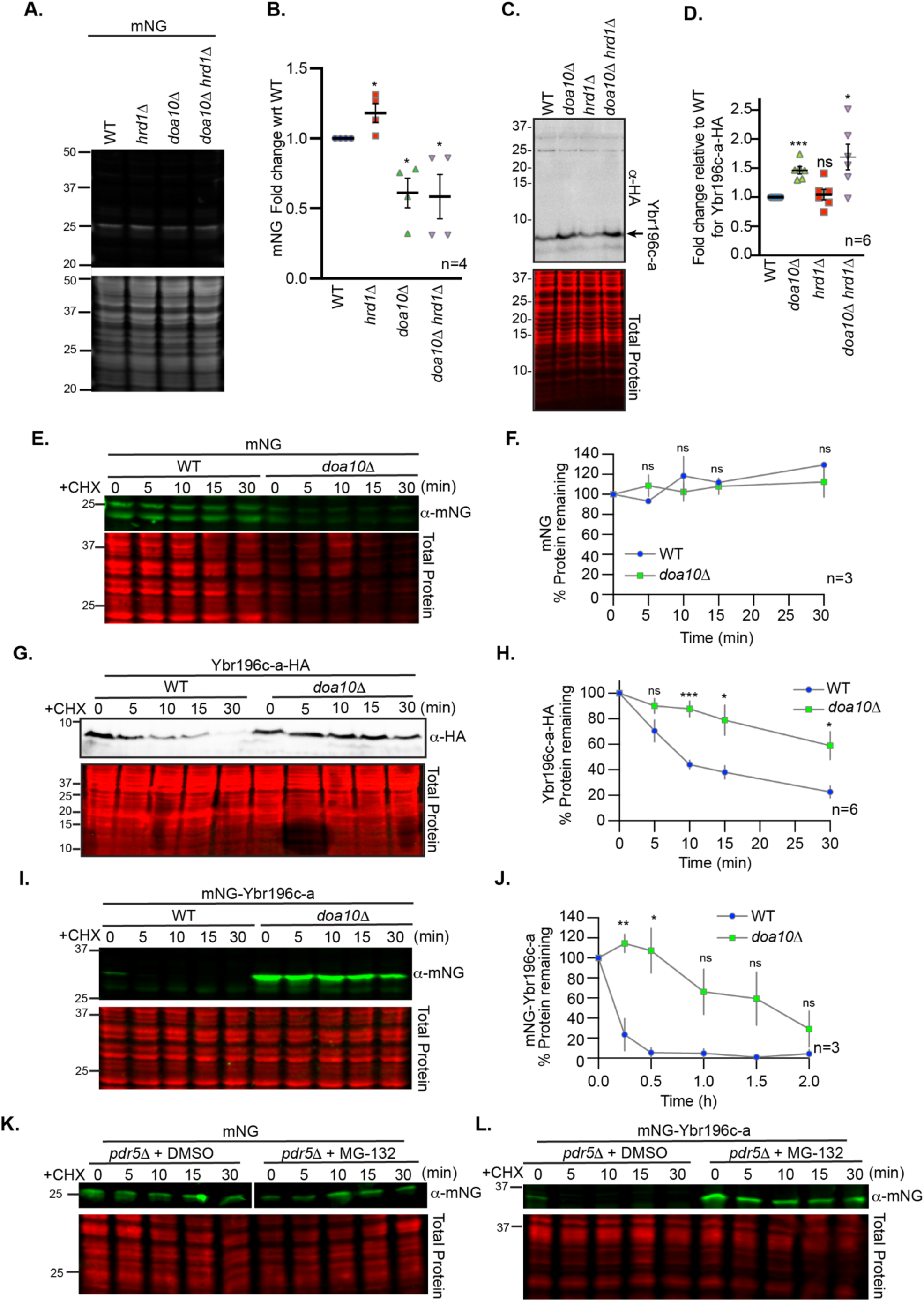
mNG stability is not dependent upon Doa10 or the proteasome, but N-terminally, mNG-tagged or C-terminally, HA-tagged Ybr196c-a stability remains dependent on both these factors. (accompanies Figure 5) **A** and **C.** Representative immunoblots of protein extracts made for **A,** free mNG, and **C**, C-terminally HA-tagged Ybr196c-a in cells with the indicated gene deletions. **B** and **D.** The relative abundance of **B,** free mNG, or **D**, Ybr196c-a-HA from biological replicates (n) of the immunoblots in **A** and **C**, respectively. A Student’s t-test comparing the WT to the Ub ligase deletions was used to assess statistical differences (ns: not significant; *:p < 0.05; ***:p < 0.0005). **E, G,** and **I.** Representative immunoblots of protein extracts expressing **E**, free mNG, **G**, Ybr196c-a-HA, and **I,** mNG-Ybr196c-a in WT or *doa10*Δ cells post cycloheximide (CHX) addition for the times indicated (in minutes) are shown. **F, H,** and **J.** Quantification of the percent of **F**, free mNG, **H**, Ybr196-c-a-HA, or **J**, mNG-Ybr196c-a remaining post-CHX addition is plotted. The symbols represent the mean of the biological replicates (n), and the error bars show the standard error of the mean. A Student’s t-test comparing the WT to the *doa10*Δ cells at each time point was used to assess statistical differences (ns: not significant; *:p < 0.05; ***:p < 0.0005). **K-L.** Representative immunoblots of protein extracts expressing **K**, free mNG or **L**, mNG-Ybr196c-a in *pdr5*Δ cells that were untreated (+DMSO) or treated with the proteasomal inhibitor MG-132 post-CHX addition are shown. Deletion of Pdr5 allows for better retention of MG-132 in cells.

**Figure S7.**
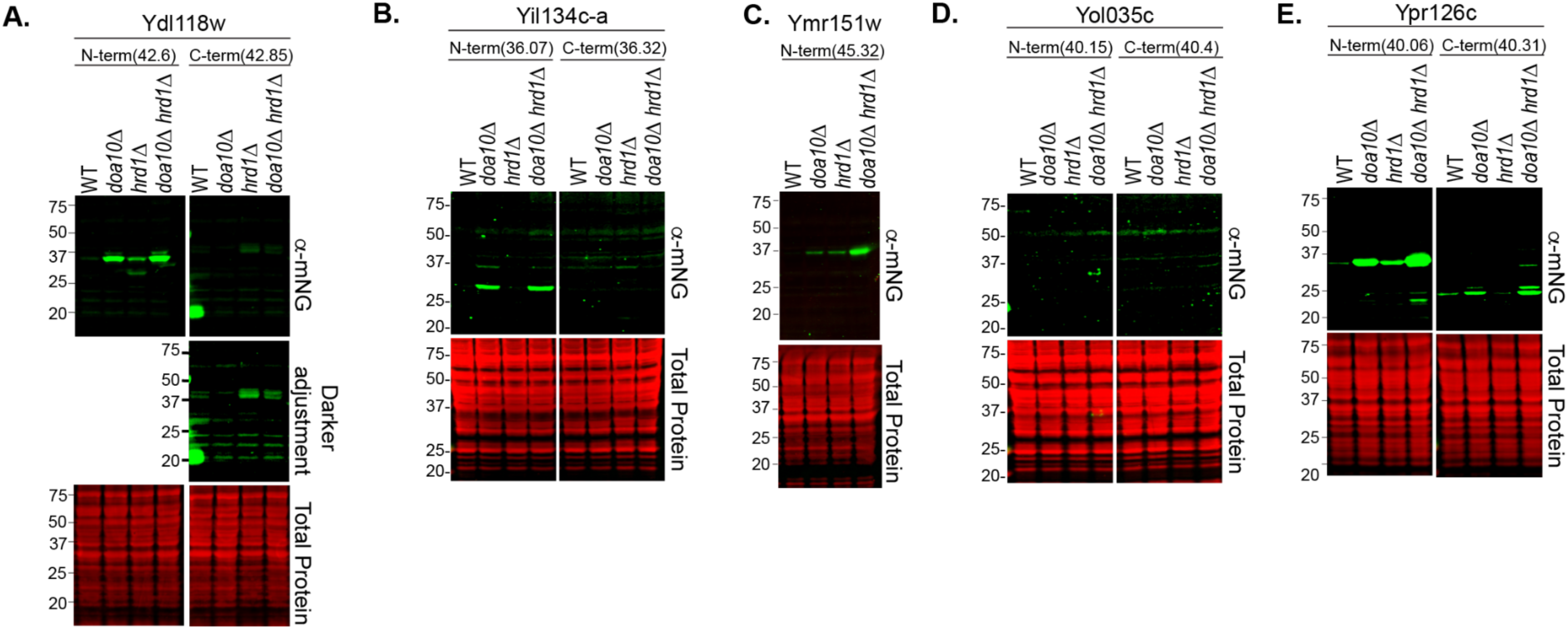
Most ER-localized DNPs are stabilized by loss of E3 Ub ligases Doa10 and/or Hrd1. (accompanies Figure 5) **A-E.** Representative immunoblots of protein extracts made for each ER-localized DNP expressing the DNP fused N or C-terminally with mNG in the indicated cells are shown, as was done for Ybr196c-a in Figure 5A. Expected sizes (MW) of the DNP fusions that are N-terminally (N-term) or C-terminally (C-term)-tagged with mNG are listed in parenthesis below each DNP name.

**Figure S8.**
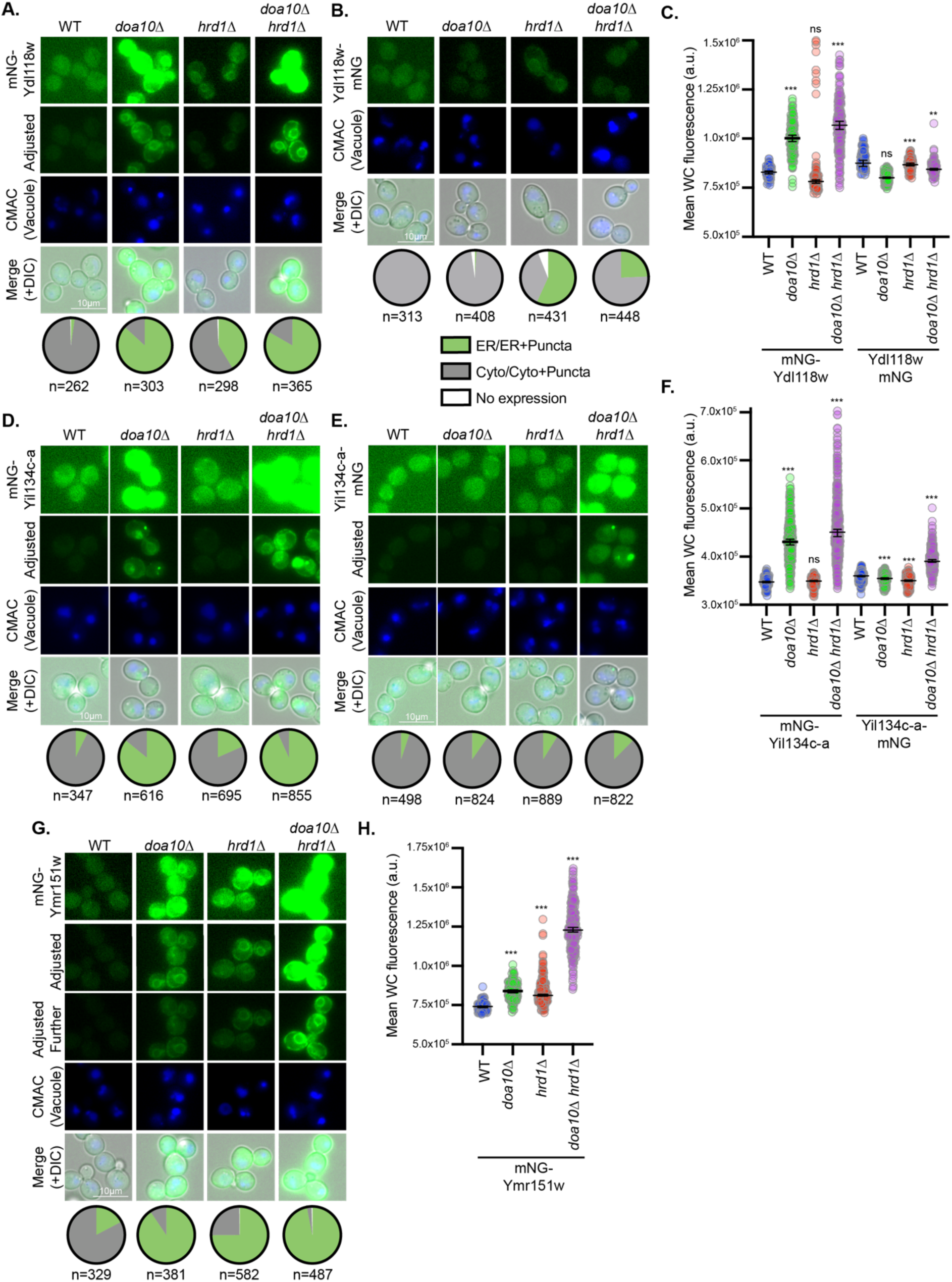
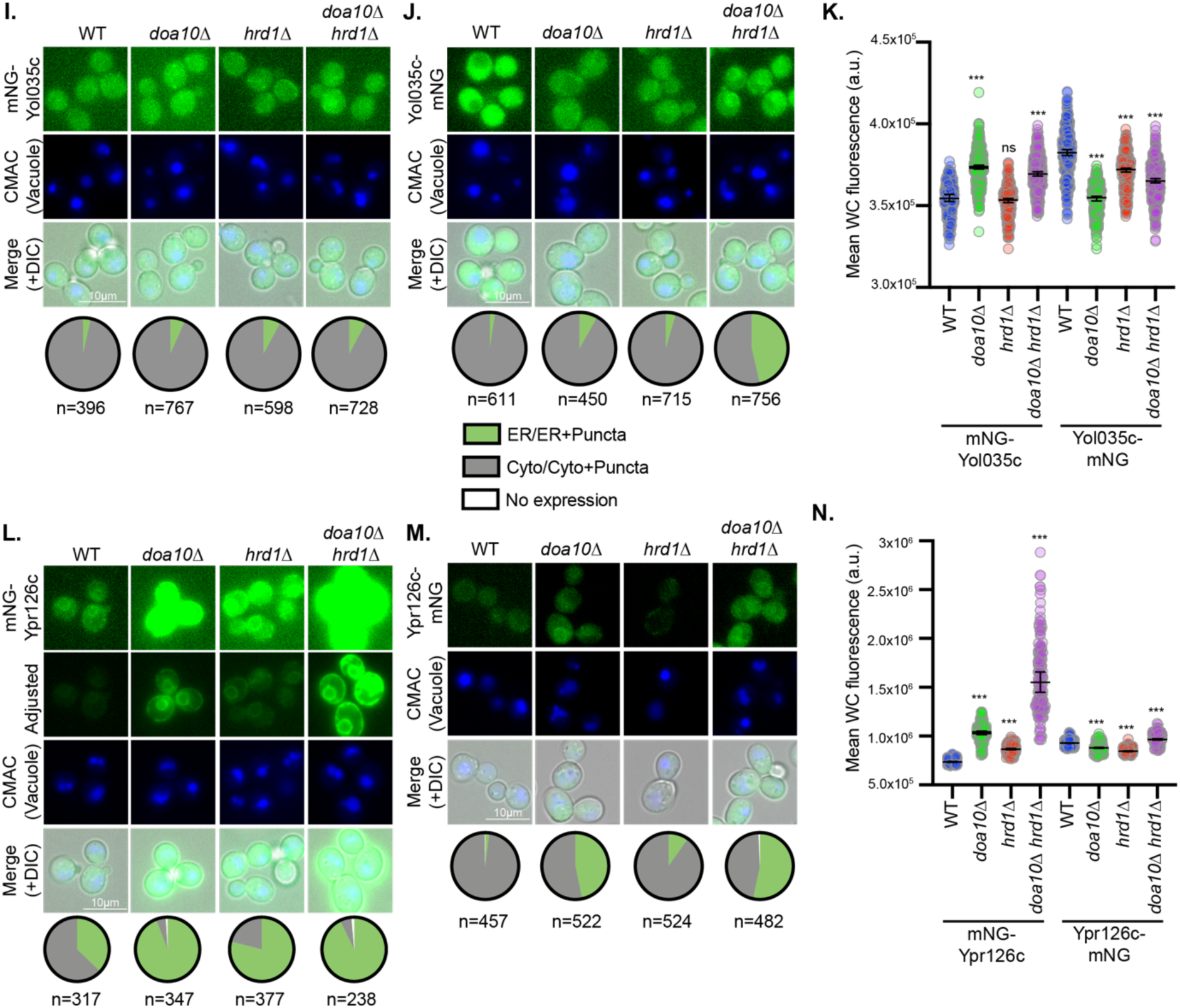
ER-localized DNPs increase in abundance when ER-resident Ub ligases are lost. (accompanies Figure 6) Representative fluorescence microscopy of N- or C-terminally tagged DNPs with mNG are shown for **A-B**, Ydl118w; **D-E,** Yil134c-a **G,** Ymr151w, **I-J,** Yol035c, and **L-M,** Ypr126c in the cells with the indicated genotypes. These are similar to the data shown for Ybr196c-a in Figure 6A and 6B. The distribution of fluorescence observed across the cell populations is indicated in the pie chart below each diagram (n=number of cells assessed). Cells are stained with CMAC blue to mark vacuoles and imaged via DIC, which is shown in the merge. **C, F, H, K,** and **N.** Quantification of the whole cell (WC) fluorescence intensities from individual cells (n=130-479) determined using the NIS.*ai* and GA3 software (see methods), for the cells shown in panels **A-B**, **D-E**, **G**, **I-J,** and **L-M,** respectively. A one-way ANOVA with Dunnett’s multiple comparisons test was performed relative to the fluorescence in WT cells to identify statistically significant changes (p-value < 0.0005 = ***).

**Figure S9.**
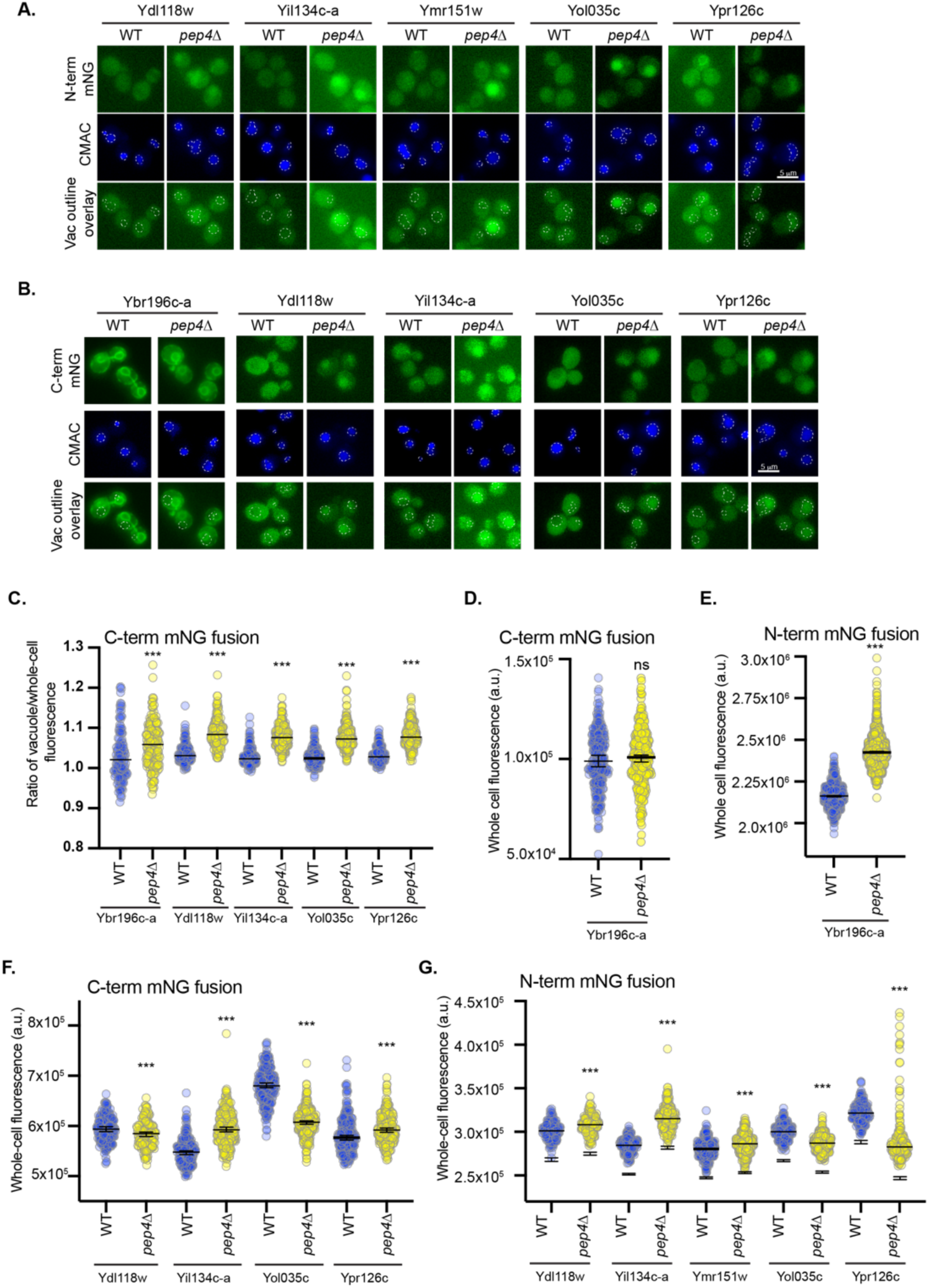
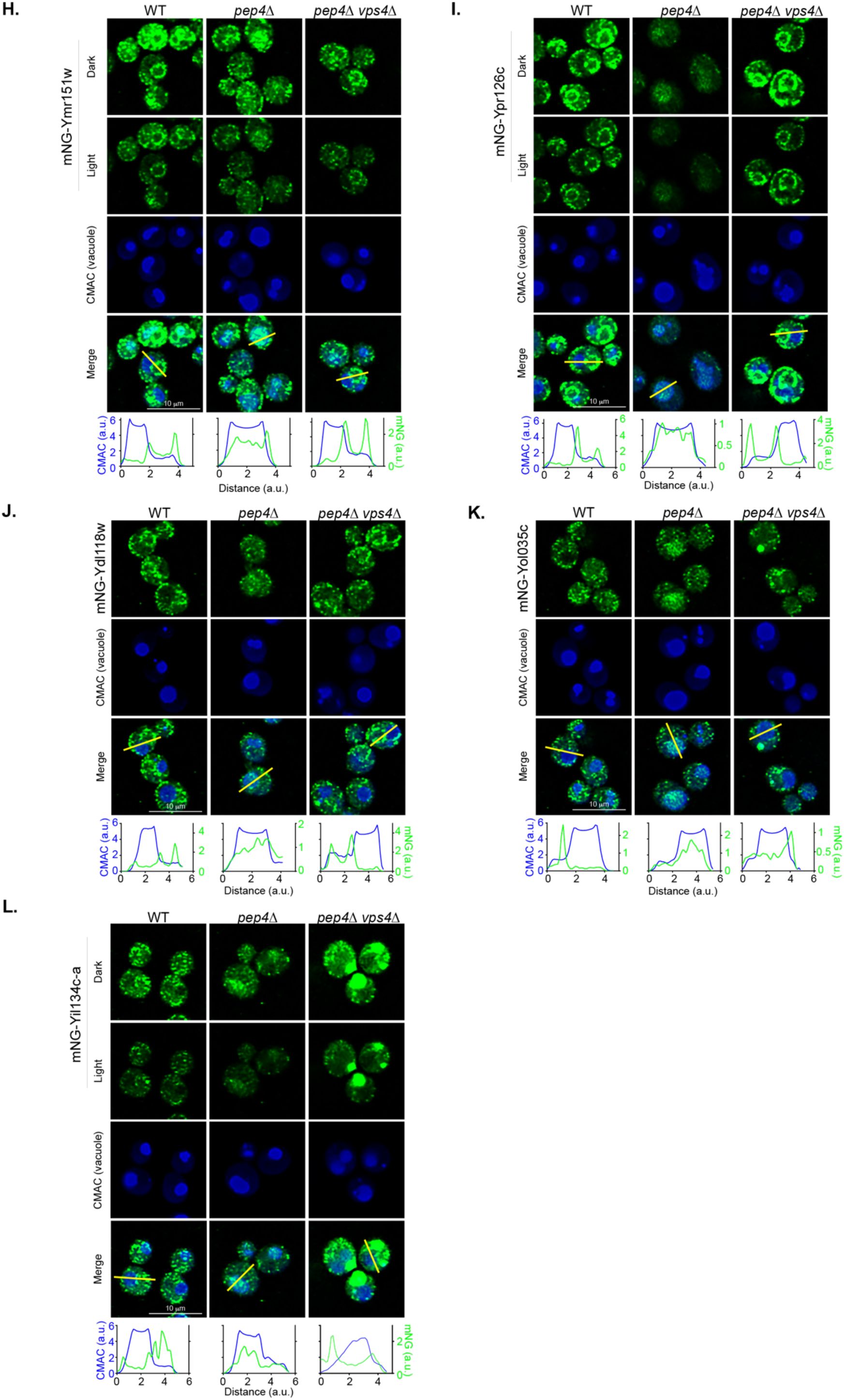
ER-localized DNPs are stabilized by loss of vacuolar proteases and depend on ESCRT III to reach the vacuolar lumen. (accompanies Figure 7) **A-B.** Fluorescence micrographs of cells expressing N-terminal in panel **A** or C-terminal mNG-tagged DNPs in panel **B** in WT or *pep4*Δ cells are shown. Vacuoles are stained with CMAC blue. Vacuoles are indicated on the green channel images using a white dashed line. **C.** Scatter plot graphing the ratio of the vacuolar fluorescence over the whole cell fluorescence for the cells imaged in panel **B** is provided. **D-G.** The scatter plots indicate the whole-cell fluorescence intensities, as determined using NIS.*ai* and GA3 software, for imaging shown in **B** (panel **D** and **F**), Figure 7A (panel **E**), and **A** (panel **G**). The median fluorescence intensity and 95% confidence interval are represented by the horizontal line. **C-G.** A Mann-Whitney two-sided test assessed statistical difference between the WT and *pep4*Δ cell populations (n=136-1294; p < 0.0005 = ***). **H-L.** Fluorescence micrographs of cells expressing mNG-DNPs in cells of the indicated genetic background are shown. Vacuoles are shown using CMAC blue. Plots of the fluorescence intensity from the blue and green channels along a line scan that runs through the vacuole (indicated in yellow on merge) are shown below each set of images. Overlapping maximum peaks of blue and green lines indicate mNG presence in the vacuolar lumen.

